# Asymmetric cortical mechanics shape epithelial cell geometry in organoids

**DOI:** 10.1101/2024.08.22.609097

**Authors:** Tristan Guyomar, Linjie Lu, Tetsuya Hiraiwa, Daniel Riveline

## Abstract

Cell cortex is a thin sheet of actin cytoskeleton spanning cell boundaries with rich out-of-equilibrium dynamics. A theoretical description of the cortex as an active fluid enables to capture cell shapes dynamics in 3D epithelial tissues. However models integrated with calibration of parameters and quantitative experiments are lacking so far. Here we report that cells in organoids and in cysts have conserved apico-basal-lateral asymmetric compositions in actin and in myosin, and we quantify their densities and mechanical properties. This allows to calibrate a new model coupling active fluids with a phase field which reproduces the main features of cell shapes. To test our approach, we successfully predict changes in cell shapes by modulating actin polymerisation, myosin activity, and adhesion in experiments and in simulations. Our study shows how active fluid theory integrated with experiments can determine cell shapes in epithelial tissues.

## I. Introduction

Cells and tissues are shaped by the acto-myosin cytoskeleton [1]. This ~200 nm thin layer represents a gel spanning each cell border. Its dynamics offers rich out-of-equilibrium spatio-temporal patterns during local protrusion [2–5], motility [6–9], or division [10, 11]. Its selforganisation properties have shown their relevance in a variety of conditions *in vitro* [12, 13] *and in vivo* [14–16]. This level of achievement results from a progress in the experimental physics of actin and acto-myosin *in vitro* associated with the development of active fluid physics [17]. Actin gels were shown to promote bead motion through polymerisation *in vitro* [18–20] and acto-myosin gels were reported to undergo contraction [21]. These phenomena were compared to the theory of active gel revisiting the Navier-Stokes equation by adding a new term taking into account the stress generation internally driven by the acto-myosin cortex [17, 21–24]. Progress over the past decades has enabled theoretical predictions and their relevance mainly in single cell often in 2D [10, 25, 26].

However, at this stage, tests of this cytoskeleton/active gel approach are needed in more physiological situations. Cells grow in 3D in *in vivo* environments. Also cell cortex requires specific measurements to evaluate its physical properties in order to associate theoretical parameters with actual physical characteristics in experiments. These limitations to extrapolate results obtained so far call for new approaches combining control of 3D tissue growth with quantitative experiments allowing to calibrate active fluid based simulations.

In this context, epithelial cysts represent interesting systems. They are composed of cell monolayers surrounding a fluid filled cavity called the *lumen* [27–29]. Cysts are also reported as the basic structure at the origin of organs [27], ranging from the formation of embryos to the generation of pancreatic ducts for example [28, 31]. With their *in vitro* controls and their relevance for organogenesis with properties shared with organoids, cysts appear as ideal systems to address this interplay between the physics of acto-myosin cortex and its theoretical description with active fluid formalism.

In this Article, we quantitatively investigate the morphological and mechano-chemical features of cell shape in epithelial cysts. Using Madin–Darby Canine Kidney (MDCK) cysts as a model system, we quantify the spatial organization of the acto-myosin cortex, measure its turnover dynamics and mechanical properties, and characterize the resulting cell geometries under a range of perturbations. To determine whether these experimentally measured cortical asymmetries are sufficient to account for epithelial morphology, we develop a multicellular active-gel model coupled to a phase-field description of cells, the extracellular matrix and the pressurized lumen. Using a single calibrated mechanical framework, the simulations reproduce the morphological trends induced by perturbations of actin dynamics, myosin activity and cell–cell adhesion, supporting the proposed physical mechanism underlying epithelial cyst morphogenesis.

## II. Materials and Methods

### A. Experimental system

#### 1. Cell culture

Madin-Darby Canine Kidney (MDCK) II cells were cultured using Gibco Minimum Essential Media (MEM) supplemented with 5% Fetal Bovine Serum (FBS). At 60-80% of confluency (every 2-3 days), cells are replated at a seeding density of 4.10^3^ cells.cm^−2^. The MDCK cell lines used in this work with fluorescently labeled proteins are listed in SM[32].

#### 2. Sample preparation

We obtained MDCK cysts lying within the same focal plane using a protocol adapted from Refs. [33, 34]. Coverslips were activated by O_2_ plasma and coated with a 100% Matrigel solution (Corning BV: 356231). Single cells were deposited at a density of 15,000 cells/cm^2^ and were covered next by another layer of 10 *µ*l 100% Matrigel. Results after 5 days of culture are presented in Fig.1 and at 3, 5 and 7 days of culture in Appendix Fig.3. For mES epiblasts and pancreatic spheres, cells were centrifuged into micro-cavities and their diameters were adapted to contain a single cells as described in [36].

For quantification of cell shape and co-labeling of actin, myosin and E-cadherin distributions, we relied mostly on fixed samples that allow higher quality imaging. For perturbation experiments, we used live samples and stable cell lines reporting for actin or myosin.

#### 3. Immunostainings

For immunofluorescence microscopy, Myosin-Regulatory-Light-Chain (MRLC) was stained with a rabbit anti-MRLC antibody (#3672; 1:500; Cell Signaling Techonologies), F-actin with phalloidin (A12379; 1:400; ThermoFisher) and E-cadherin with a rat anti-E-cadherin antibody (Ab11512; 1:500; Abcam) as described in [34].

#### 4. Microscopy

Fixed samples imaging and time-lapse imaging were performed with a Leica DMi8 confocal microscope equiped with a Yokogawa CSU W1 spinning disk and with an OrcaFlash 4.0 camera. We used a HC PL APO 63x glycerol objective (Leica - numerical aperture (N.A. = 1.3), working distance (W.D. = 0.3 mm)). For fixed samples, we acquired 3D stacks with a z-step of 0.3 *µ*m and x-y resolution of 0.103 *µ*m. When using live samples, an incubation chamber was maintained at 37°C temperature and 5% CO_2_.

Fluorescence Recovery After Photobleaching (FRAP) experiments were conducted on this spinning disk microscope using the FRAP module on the Metamorph software (iLas2). Details of this experiment and its calibration can be found in SM [32].

Laser ablation experiments were performed using a TCS SP5 inverted microscope setup under temperature controlled environment with a Z-galvo stage. The microscope was controlled using the Leica LAS AF software with the FRAP Wizard module. This allowed to control a pulsed MP laser for ablation (Coherent Chameleon Ultra II, 800 nm, 80 MHz, 4 MW). For the laser ablation experiments, we used the HC PL APO 63x oil lambda blue objective (N.A. = 1.4 and W.D. = 0.1 mm) with a numerical zoom factor of 3. Details of this experiment and its calibration can be found in SM [32]. For all microscopy data further analysed in this work, additional information regarding image analysis can be found in Appendix 2 and SM [32].

#### 5. Drug treatments

To investigate the role of actin polymerisation in shaping cells and cysts, we used 200nM of an inhibitor of actin polymerisation latrunculin A (LatA), (Sigma Aldrich #L5163) and an activator of actin polymerisation C8N6 BPA polyamine (C8BPA) [35] on 5-day-old cysts. Timelapses of samples were acquired under the microscope and the medium was then replaced by a medium containing 200 nM of LatA or 100 *µ*M of C8BPA. In the case of LatA, following 1 hour of incubation, samples were washed 3 times with fresh medium to test the successful reversibility of drug treatement. In the case of C8BPA, the sample was treated and imaged for up to 12 hours following the drug treatment. Controls for both these experiments are DMSO (dilution*<* 1%) treated cysts.

To investigate the role of cell contractility in shaping cells and cysts (Appendix Fig.8), we incubated 5-day-old MDCK cysts with *c* = 10 *µ*M of myosin inhibitor Blebbistatin (Sigma B0560) and used DMSO treated cysts as control [34].

To investigate the role of cell-cell adhesion in shaping cells and cysts (Appendix Fig.9), we used 5 mM EDTA and E-cadherin 10 µg/ml antibody [36–38].

#### 6. Statistical tests

When violin plots are used to visualize statistical distributions, the central line corresponds to the mean, the extrema lines correspond to the minimum and maximum of the distribution. All the p-values correspond to one tailed tests except for Appendix Fig.6 and Supp. Fig. 5 (c-f) where one tailed tests of related samples of scores are used. Values of *p >* 0.05 are deemed non-significant. We indicate in each figure caption the precise threshold for defining the quality of the tests (*, **, ***, n.s).

### B. Numerical simulations

#### 1. Multicellular phase-field model

We constructed our mathematical model based on the phase field (PF) approach. Since our preliminary analysis revealed that the features observed in a 2D section can sufficiently represent those in 3D (Appendix 1(a), Appendix Fig.1), we formulated our model in 2D.

We simplified cysts into three components: cells, lumen and ECM. Their geometries are described by using PF variables *ϕ*_*i*_(***r***, *t*) (*i* = 1, …, *m*; one for each cell), *ℓ*(***r***, *t*) and *e*(***r***, *t*), respectively. The regions where each PF variable takes 1 and 0 represent the inside and outside of the corresponding phase, respectively (Fig.2).

**FIG. 1.**
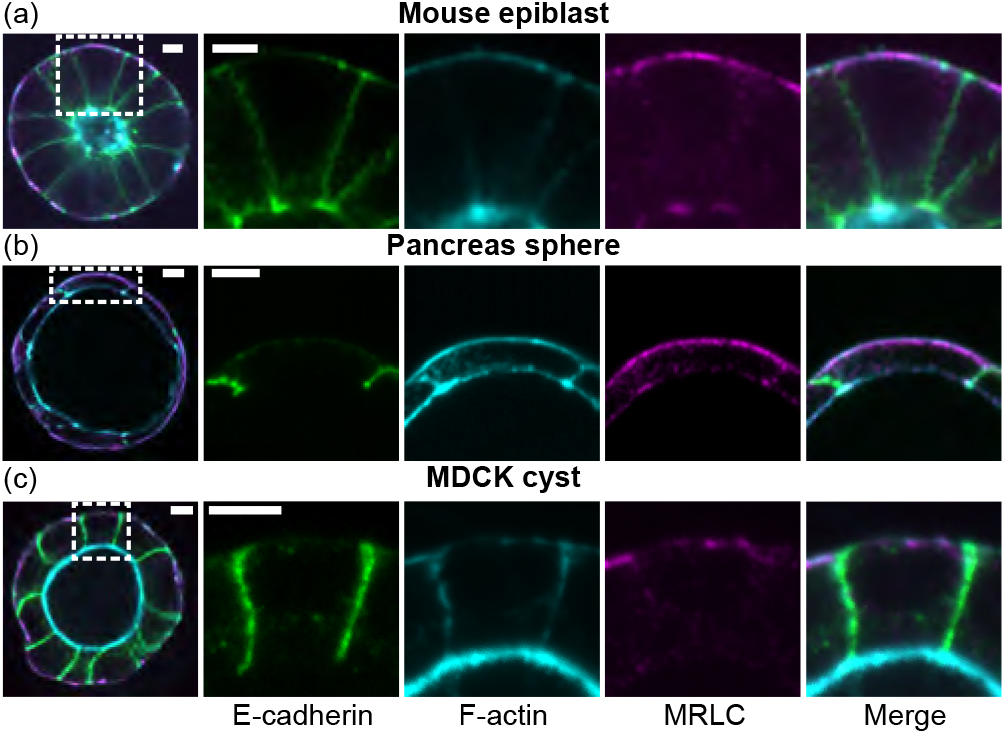
Distribution of acto-myosin in 5-day-old mES organoids, pancreatic sphere organoids and MDCK cysts. Snapshots of a mouse epiblast organoid derived from mES cells (a), of a mouse pancreatic sphere (b) and of a MDCK cyst 2D cut and zoom in, fixed and stained for E-cadherin (green), F-actin (cyan), myosin (magenta). Scale bars, 5 µm.

**FIG. 2.**
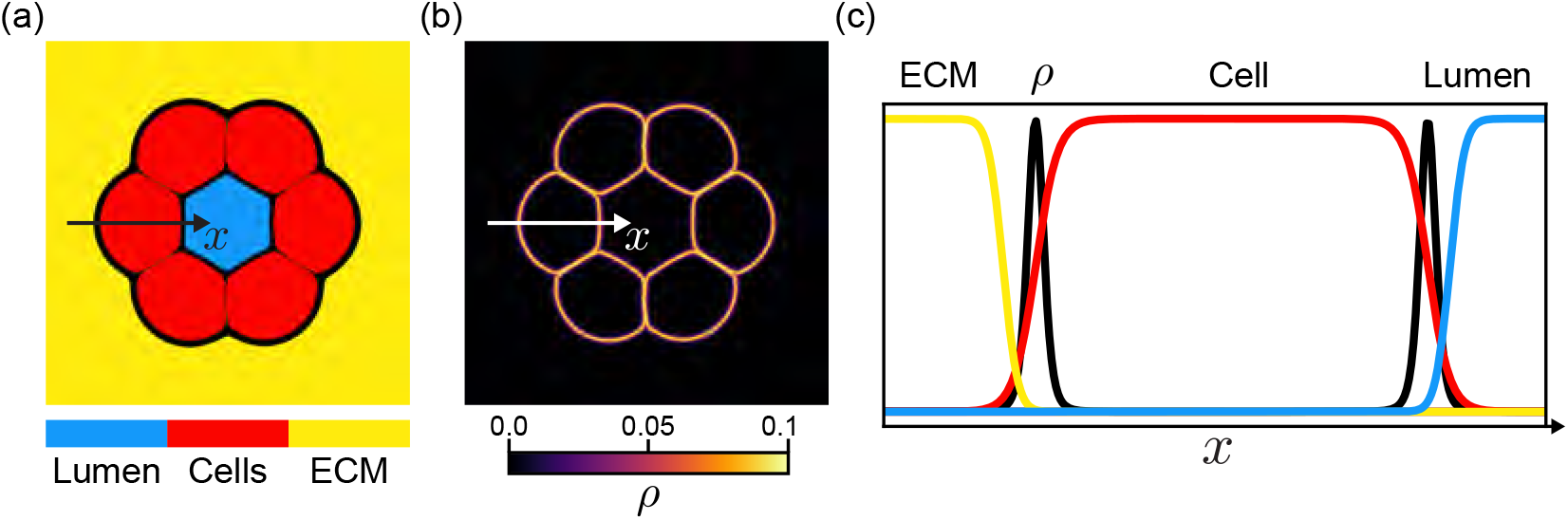
Schematic representation of the multicellular phase-field model. (a) Phase field model of cells, lumens, and ECM in 2D. (b) Active gel density ρ representing the active cortex of each cell. (c) Cross-sectional profiles of the phase fields (lumen, cell, ECM) and the active gel density ρ along the line in (a) and (b). Red, cyan, and yellow colors correspond to the phase fields of cells, lumen, and ECM, respectively, for both (a) and (c).

Each PF variable obeys mainly the standard multicellular PF model [36, 39–41], under the assumptions: (i) all cells have identical optimum cell size, cell-cell adhesion, cell-ECM adhesion and excluded volume; (ii) dynamics of lumen volume are driven by osmotic pressure differences; and (iii) the ECM is described as an elastic material (all details on the model is accessible in SM [32] as well as complementary information regarding numerical implementation, a discussion of the choice of phase-field model and sensitivity analysis).

#### 2. Coupling a cortical active gel model with a cellular phase field

We applied the resharpening method proposed in [42, 43] which suppresses the inherent surface tension emerging from basic construction principle of the phase field model, to assure that the cellular surface tension comes solely from active gel and that the surface tension of lumen and ECM can be well controlled.

We defined for each cell a cortical gel confined in the vicinity of the cellular interface, set by *ϕ*_*i*_(*r*) = 1*/*2, which is modeled as an active gel of density field *ρ*_*i*_(***r***, *t*) obeying the 2D extension of the 1D model developed in Refs.[44, 45]. Dynamics of the field variables is given in SM [32].

## III. Results

We first cultured mouse embryonic stem cells (mES) organoids [36, 46], pancreatic sphere organoids [36, 47] and MDCK cysts. After fixation and staining for F-actin, MRLC and E-cadherin, we observed that actin and myosin were asymmetrically distributed around each cell (Fig.1). Actin was thick and enriched at the apical sides whereas the myosin layer was thinner and mainly localised at the basal sides of cells. The lateral sides were essentially even in actin distributions. These anisotropies are shared among MDCK cysts and organoids presented in Fig.1.

To further quantify cell shape and actin and myosin distributions, we focused on studying 5-day-old MDCK cysts. Cysts were invariant by rotation and we could derive a typical cell shape with its actin and its myosin distributions at the cortex in the middle cross-section of each cyst (Fig.1(c), Fig.3(a)). For additional information about comparison of 2D vs 3D signal quantifications, see Appendix 1(a), Appendix Fig.1, Supp. Fig.1, Supp. Fig.2, and Video 1.

**FIG. 3.**
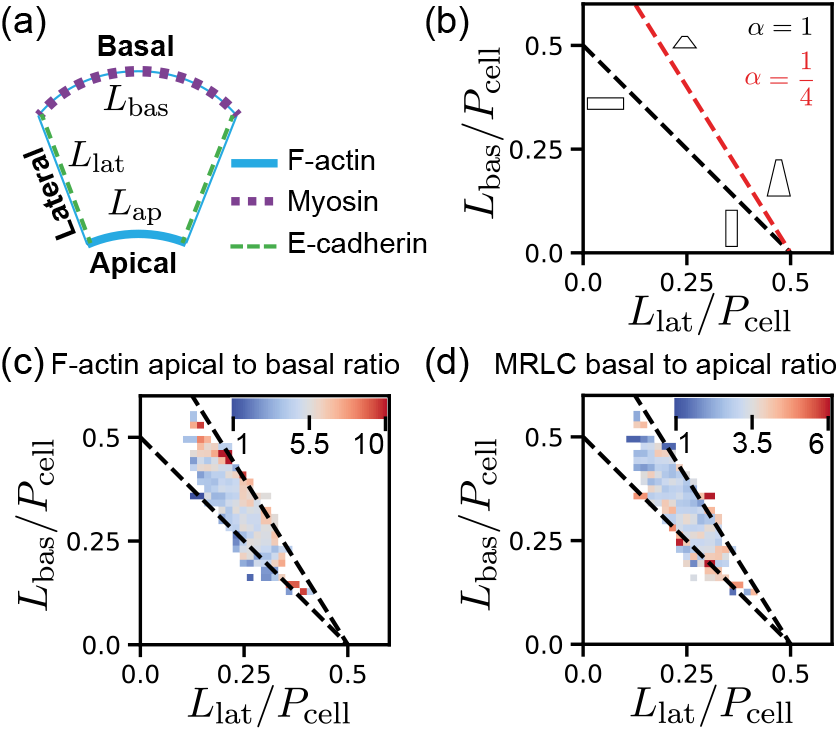
Correlation between acto-myosin cortical density and cell shape in MDCK cysts. (a) Typical mean cell description used to quantify the relevant geometrical quantities to describe the cell cortex and the cell shape. (b) Schematic description of the cell shape diagram. Oblique lines correspond to different apical to basal length ratios α as described in the main text. Moving along the line changes the aspect ratio of cells w.r.t. the lateral length. (c) F-actin apical to basal intensity ratio correlation with cell shape. (d) Myosin basal to apical intensity ratio correlation with cell shape. P_cell_, cell perimeter. For all measurements, N = 3 experiments, n = 791 cells quantified in 5-day-old MDCK cysts.

We measured that the F-actin intensity was about 5 times higher in the apical cortex than in the basal one and about 7 times higher in the apical cortex than in the lateral one (Appendix 1(c), Appendix Fig.2(b-c) and Table I). We found that the myosin intensity was approximately 3 times higher in the basal cortex than in the apical cortex and about 4 times higher in the basal cortex than in the lateral one (Appendix 1(c), Appendix Fig.2(b-c) and Table I). Such anisotropies were conserved in time for MDCK spheroids (Appendix Fig.3).

**TABLE I.**
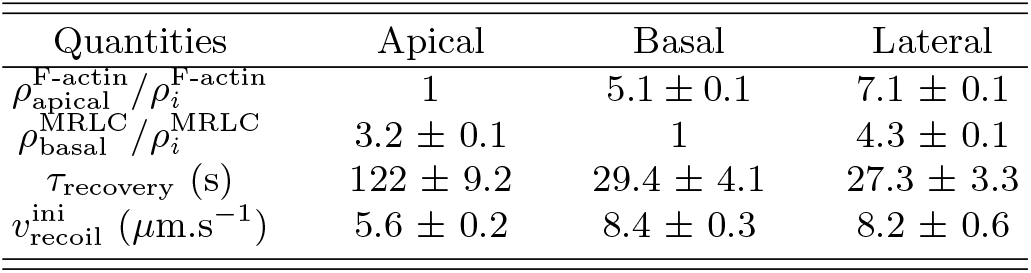
Comparison of the apical-basal-lateral measurements with intensity ratios, FRAP and laser ablation experiments. Ratios of ρ corresponds to ratios of intensities. Measurements are shown as: mean ± standard error of the mean.

Next we quantified and tested potential correlations between cell shapes and the actin and myosin concentration at the cortical layers. We represented the cell shape in a diagram that can capture most of its morphological attributes. We define *L*_ap_, *L*_bas_, *L*_lat_ as respectively the apical, basal and lateral lengths and the total cell perimeter *P* that we measured on fixed samples at day 5 of culture (Fig.3(a)). By construction, *P* = *L*_ap_ + *L*_bas_ + 2*L*_lat_. By defining the apical length as a fraction of the basal length: *L*_ap_ = *αL*_bas_, one can represent 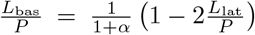 (Fig.3(b)). We used this cell shape representation to look for correlations between cell shape and intensity ratios of actin and myosin, curvatures and lumen occupancy (defined as the ratio of lumen to cyst volumes) (Fig.3, Appendix Fig.4, Appendix Fig.6, Appendix 1(d-e)). We noted a strong correlation between F-actin apical to basal ratio and trapezoidal shaped cells (Fig.3(c) Appendix Fig.5(a-c)). On the other hand, MRLC basal to apical ratio was weakly negatively correlated to a trapezoidal cell shape (Appendix Fig.5(d-f)). We verified that these correlations measured for MDCK cysts at day 5 were also present for cysts cultured at day 3 and day 7 (Appendix 1(e), Appendix Fig.3, Appendix Fig.4, Appendix Fig.5).

Localisation alone was not yet informative about the mechano-chemical parameters of these active gels. To interrogate their characteristics, we used the Fluorescence Recovery After Photobleaching (FRAP) and laser ablation of apical, basal and lateral sides with MDCK cysts cultured from MDCK cell lines stably expressing fluorescent Green Fluorescent Protein (GFP) fused proteins, actin-GFP or myosin regulatory light chain-GFP (MRLC-GFP), see SM [32]). Results are given in Fig.4 and in Videos 2-5 and experimental procedures are detailed in Appendix 2. Fig.4(d-f) show the measurements obtained through the fitting procedure (Appendix 2) that we applied to extract the characteristic time of the recovery after photobleaching that we performed at apical, basal and lateral cortices. We extracted three experimentally relevant parameters: mobile fraction, *τ*_delivery_ and *τ*_diffusion_. FRAP experiments revealed that the mobile fraction of the apical cortex was lower than the basal and lateral ones. The characteristic time of diffusion (orders of seconds, similar to reported values [48, 49]) of G-actin monomers was small compared with the characteristic time associated to transport and assembly of F-actin filaments and network (orders of tens of seconds) (Fig.4(g-i)). Interestingly, the time of transport was different by a factor of 4 between apical and basal sides, which substantiates again the asymmetry between sides.

**FIG. 4.**
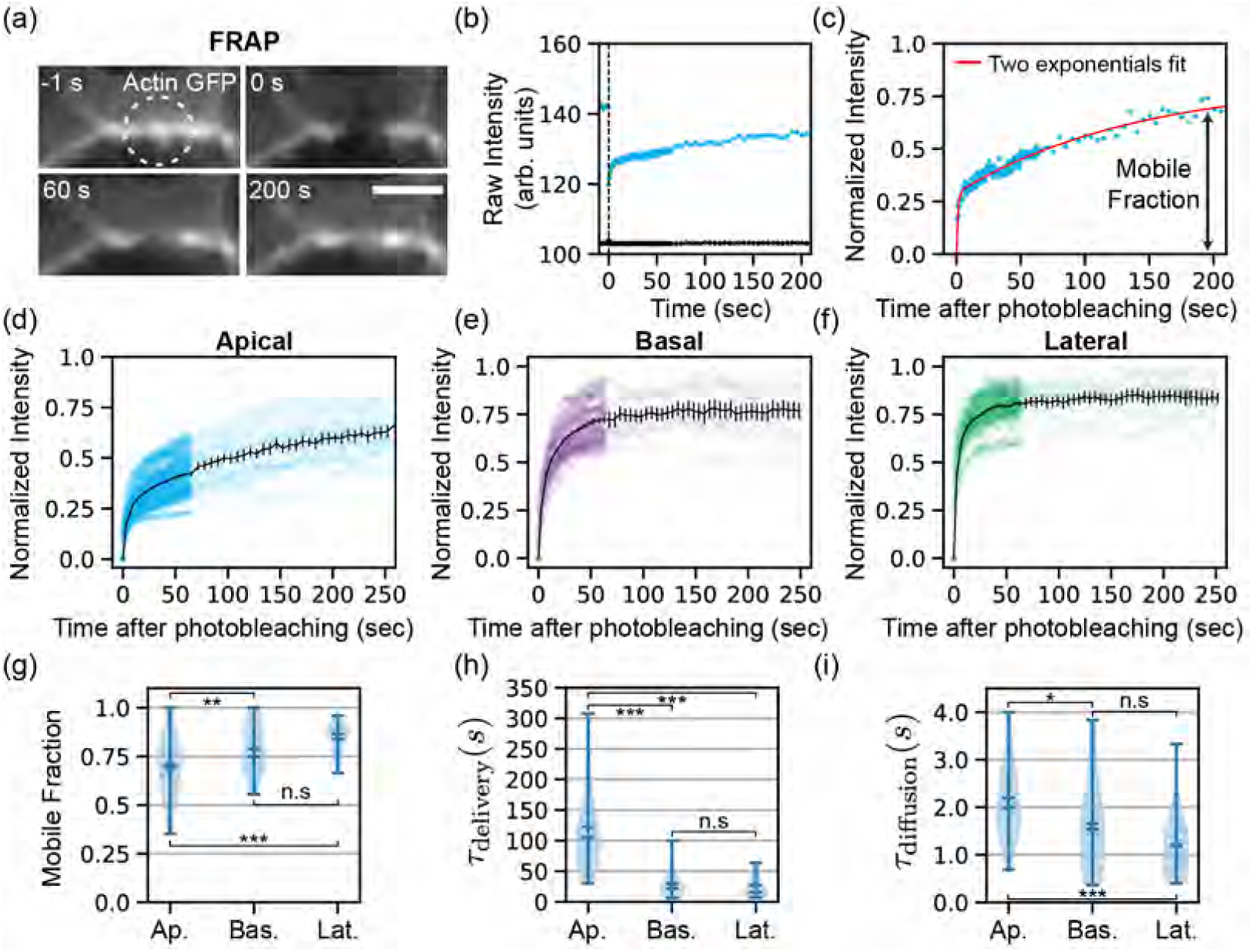
Quantification of the actin gel turnover properties at different locations of the cell cortex in day 5 MDCK cysts by FRAP. (a) Snapshots of FRAP experiments performed on the apical cortex of MDCK cysts labelled for actin-GFP (grey) (Video 2). (b) Raw actin intensity as a function of time after photobleaching (raw intensity (blue) background (black)). (c) Normalised actin intensity as a function of time of the raw actin intensity shown in (b). The red line corresponds to the fit with a sum of two exponentials. Normalised actin intensity as a function of time after photobleaching of the apical cortices (d), basal cortices (e) and lateral cortices (f). Black curves represent the average normalised intensity recovery after photobleaching. Error bars: 95 % confidence interval of the mean. Measurements of (g) mobile fraction, (h) τ_delivery_ and (i) τ_diffusion_ at apical, basal and lateral cortices. N = 3 experiments, n = 60 cells (apical), n = 30 cells (basal and lateral). Statistical tests: *: p < 0.05, **: p < 0.01, ***: p < 0.001, n.s: p > 0.05. Scale bars : 5 µm.

Finally, we determined that both sides were contractile based on laser ablation experiments and initial recoil velocity, 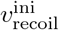, measurements (Fig.5 and Supp. Fig. 4). These experiments allowed to extract mechano-chemical parameters summarized in Table I.

**FIG. 5.**
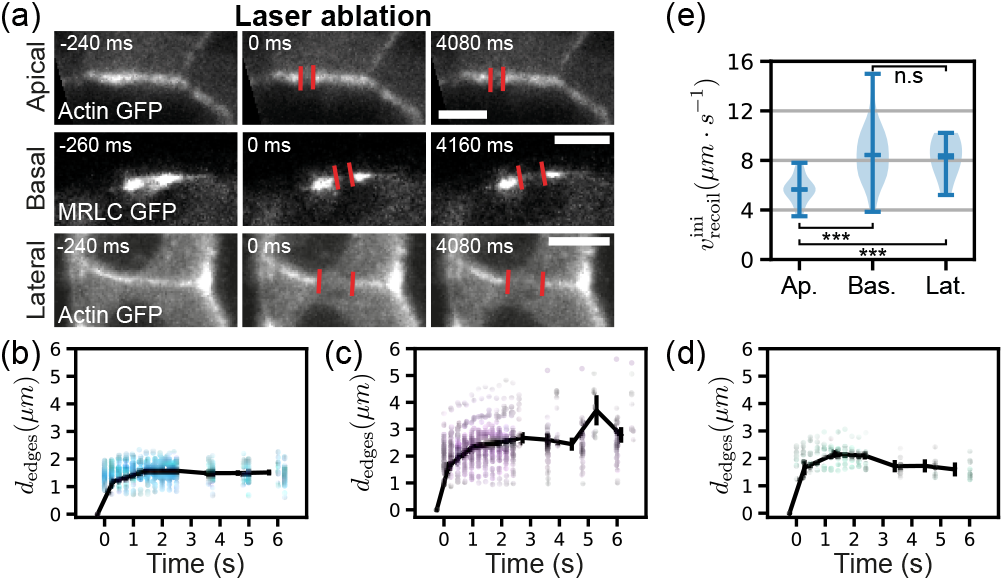
Quantification of the acto-myosin tensile properties at different locations of the cell cortex in day 5 MDCK cysts by cortical laser ablation. (a) Snapshots of laser ablation of the cortex performed on the apical, basal and lateral cortex of MDCK cysts labelled for actin-GFP (grey) apical and lateral) or MRLC-GFP (grey - basal) (Videos 3-5). (b)-(c)-(d) Distance between the edge of the ablated cortex d_edges_ as a function of time after laser ablation for apical, basal and lateral cortices. Error bars: 95 % confidence interval of the mean. (e) Initial recoil velocity for apical, basal and lateral sides measured from the initial distance created after laser ablation. N = 4 experiments, n = 43 cells (apical), n = 45 cells (basal), N = 3 experiments, n = 8 cells (lateral). Statistical tests: *: p < 0.05, **: p < 0.01, ***: p < 0.001, n.s: p > 0.05. Scale bars : 5 µm.

Next, we theoretically tested our hypothesis that the asymmetry in cortex distribution is a key factor to determine cell shape in a cyst, through numerical simulations based on multicellular phase-field (PF) model [36, 39–41] coupled to active gel equations [44, 45]. All detailed equations, numerical schemes, choice and sensitivity analysis of parameters of the model can be found in SM [32].

The geometry of cells is described by PF variables *ϕ*(***r***, *t*) (Fig.2), defined for each cell, and the cortical gel is confined in the interface of cellular PF by the coupling free energy:

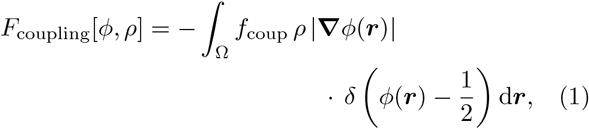

with *f*_coup_ controlling the strength of coupling and Ω the simulation domain area. This free energy term adds a new term to the cellular-PF dynamics (see Eq. (S27) of SM [32]). This coupling term induces a global contractility which counteracts the volume control and adhesion energies already set in the cell shape dynamics. *ρ*(*r, t*) obeys the conservation equation with the source of cortical gel at the cellular-PF interface:

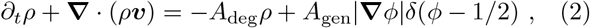

where the first and second terms on the right-hand side are the degeneration and source at the cell interface, respectively.

*A*_deg_ and *A*_gen_ represent the turnover rate and production speed, respectively. *v*_*α*_(*r, t*) is the gel velocity field and given by the force balance: *∂*_*β*_*σ*_*αβ*_ = *γv*_*α*_, where *σ*_*αβ*_ is the total stress tensor, *γ* is a friction coefficient and *α, β* = *x, y*. We write the stress tensor as *σ*_*αβ*_ = *σ*_viscous_ + *σ*_active_ = 2*ηv*_*αβ*_ − Π_*αβ*_(*ρ*) with the velocity gradient tensor *v*_*αβ*_ = (*∂*_*α*_*v*_*β*_ + *∂*_*β*_*v*_*α*_)*/*2 and choose to follow Ref.[44] to express Π in terms of the gel density *ρ* with *π*_*αβ*_(*ρ*) = −*a*_*αβ*_*ρ*^3^ + *b*_*αβ*_*ρ*^4^. For *a >* 0, the active gel is contractile whereas, for *a <* 0, the active gel is extensile.

We performed the numerical simulations based on the equations presented above (SM [32]). Taking advantage of the phase field formulation of the model, we defined apical, lateral and basal cortices and imposed different parameters to each of them (SM [32] and Table S2 for the complete set of parameters values, the experimental correspondance, and sensitivity analysis). We modeled a 2D cut of a cyst composed of 6 cells, and we depicted the main results in Fig.6 and in Video 6. Our first aim is to explore the possibility offered by our numerical model and to display how different asymmetries in the cell cortex properties can yield a variety of cells and cysts shapes (Fig.6(a-f), SM [32]). Our second aim is to explore the model parameter landscapes by using experimental measurements of the cell cortical properties and find a set of parameters that faithfully reproduces experimental cell and cyst morphologies (Fig.6(g), Appendix 3, Table I, SM [32]). Our third aim is to test our model by modifying some key parameters of the model and compare the changes in cysts and cells shapes both in experiments and numerical simulations (Fig.7, Appendix Fig.7-10, Appendix 4, SM [32]).

**FIG. 6.**
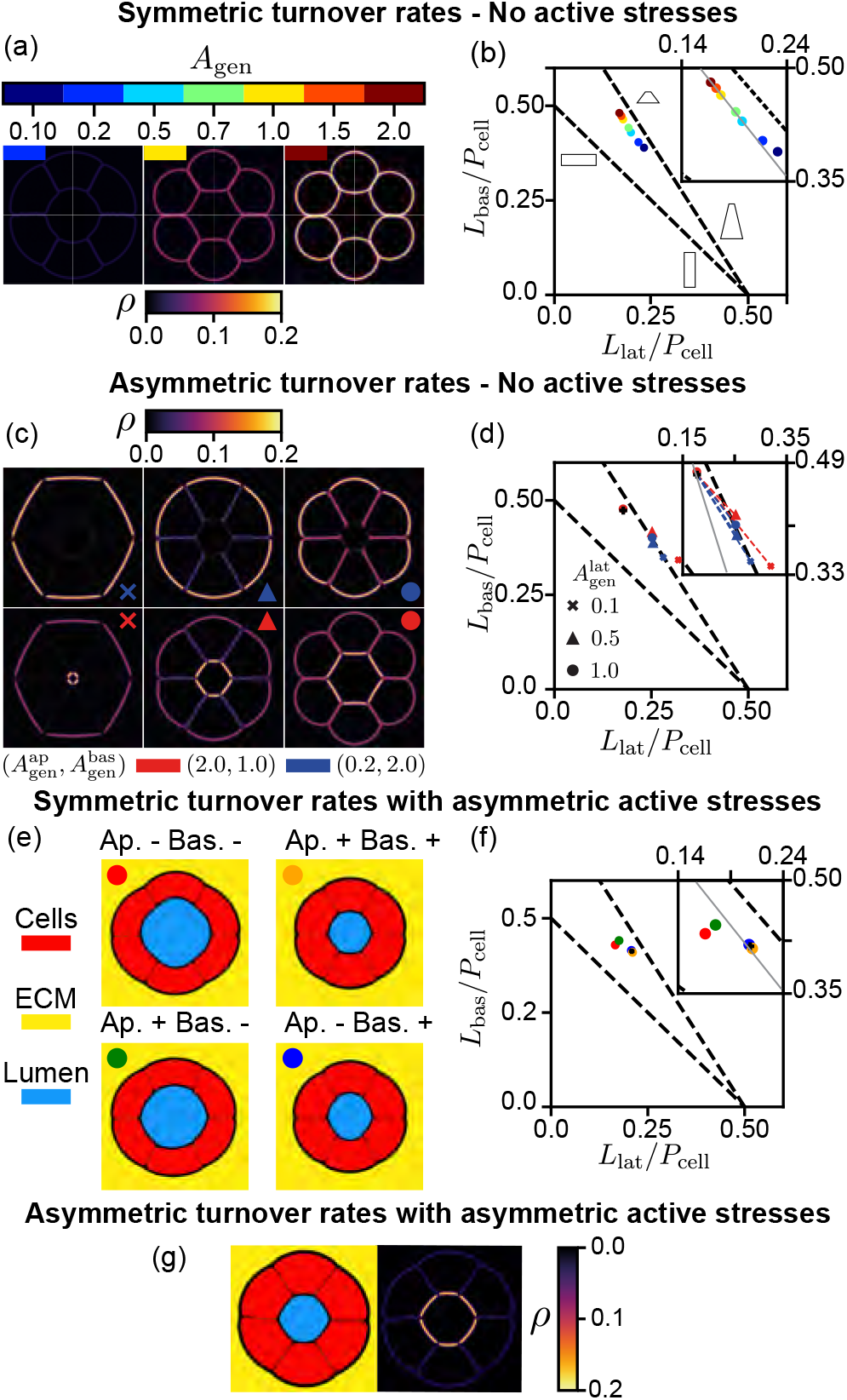
Asymmetry in active fluid cortices generates relevant cell shapes in numerical simulations. (a) Snapshots of simulated 2D cuts of cysts with symmetric apical, basal and lateral turnover rates at steady state for different turnover rate magnitudes, active gel density ρ is represented (Video 6). (b) Cell shape evolution when A_gen_ is modified globally. (c) Snapshots of simulated 2D cuts of cysts with asymmetric turnover rates. (d) Cell shape changes associated to asymmetry in turnover rates. Same symbols in (c) and (d). (e) Snapshots of simulated 2D cuts of cysts with symmetric turnover rates but asymmetric additional extensile/contractile active stresses. Cells (red), lumen (blue) and an elastic ECM (yellow) are represented. All simulations started from the same initial condition and were stopped after 300000 steps. (f) Cell shape changes associated with asymmetry in active stresses. Same symbols in (e) and (f). (g) In the case of asymmetric turnover rates, parameters were chosen to match the experimental data presented in Table I. Active gel density ρ is represented in (a), (c) and (g).

We first tested how a change in uniform surface tension would impact the cell and cyst shapes by adding no additional active stresses but setting various uniform turnover dynamics in the cell cortices (Fig.6(a-b) and SM [32]). As the turnover rate increases, so does the active gel density *ρ* and the cells became rounder. Using similar diagrams as introduced in Fig.3(b), we quantified these shape changes (Fig.6(b)) and we observed that for a given set of parameters (lumen pressure, cell-cell adhesion, ECM elasticity, total cell number) the morphology of cells evolved mainly on a straight line in the cell shape diagram as *A*_gen_ varies.

Next, we implemented asymmetric turnover rates between apical, lateral and basal cortices (Fig.6(c-d)) and SM [32]). Interestingly, asymmetries in the turnover rate distributions yielded a variety of cyst morphologies (Fig.6(a)), richer than the symmetric case (Fig.6(a)). This was further illustrated in the cell shape diagram (Fig.6(d)) where asymmetric turnover rates led to departure from the previously highlighted trajectories (Fig.6(d)). We then changed the physical origin of stress generation, by testing how the asymmetry of the nature of active stresses would impact cell shapes in cysts. We kept symmetric turnover rates and we tuned the extensile/contractile nature of apical and basal cortices (Fig.6(e), SM [32]). Strikingly, whereas the active gel density remained constant, the asymmetric nature of stresses alone led to large changes in shapes. For example, in the case of apical contractile and basal extensile (Fig.6(e-f), green point), the extensility of the basal active gel yielded less active stress and more tangentially elongated cells. Finally, we used our measurements of cell cortical mechano-chemical properties and tuned a set of parameters that yielded an example of cyst that could reproduce qualitatively MDCK cell and cyst phenotypes (Fig.6(g) and Fig.1(b) bottom) when setting asymmetric turnover rates and asymmetric active stresses (SM [32]) following the measurements presented in Fig.3, Fig.4, Fig.5 and recapitulated in Table I. This procedure is detailed in Appendix 3. While it was not possible to explore a complete part of the parameter landscape for the multicellular phase-field model coupled to an active gel model of the cellular cortex, Fig.6 illustrates that this model enables us to compare qualitatively variations in the cell shape diagrams associated with changes in active gel dynamics.

Additionally, we have shown that experiments can give valuable information that can help to narrow down parameter values to generate a numerical version of cysts that we will now use to better understand how varying individual properties affects cell and cyst morphologies.

To test predictions associated to changes in the dynamics of the gel parameters (SM [32]), we first altered specifically the polymerisation of actin. Upon incubation with the depolymerising agent latrunculin A (LatA) (M&M), we recorded changes in cell shape and the associated modifications in the actin cortical composition. After treatment, cells became flatter, apical cortices bent away from the lumen and the apical actin gel depolymerised (Fig.7(a), Appendix Fig.7(a-d) and Video 7). After washout, actin cortices were rebuilt and cell shape came back to their configuration prior drug treatment in about 10 hours (Supp. Fig. 5 and Video 8). This result confirms the trend measured in (Appendix Fig.4 and Appendix Fig.5(a-b)): higher F-actin apical/basal ratios are measured in more trapezoidal cells. We also performed the opposite experiment by incubating cysts with the actin polymerisation enhancer C8BPA [35]. Actin gels polymerised following the drug incubation and cells became more cuboidal with a tendency of the apical cortex to adopt a positive curvature w.r.t. the cell center of mass (Fig.7(c,d), Appendix Fig.7(e-h) and Video 9). Quantifications are shown in Fig.7(a-d) and further discussed in Appendix. 4(a-c). Strikingly, we could obtain shapes predicted by the associated changes in actin polymerisation in the simulation (variation of *A*_gen_ in Fig.7(e,f) and Video 10). Next, we tested additional conditions in experiments and simulations. We inhibited myosin activity by incubating cysts with myosin inhibitor Blebbistatin (see Materials and Methods (M&M)) and monitored the change in lumen occupancy between 2 hours before and 2 hours after treatment (Appendix. 4(d) and Appendix Fig.8). Incubation with Blebbistatin did not change cell shape as supported also by simulation by changing corresponding parameters (Appendix Fig.8 and sensitivity analysis in SM [32]).

**FIG. 7.**
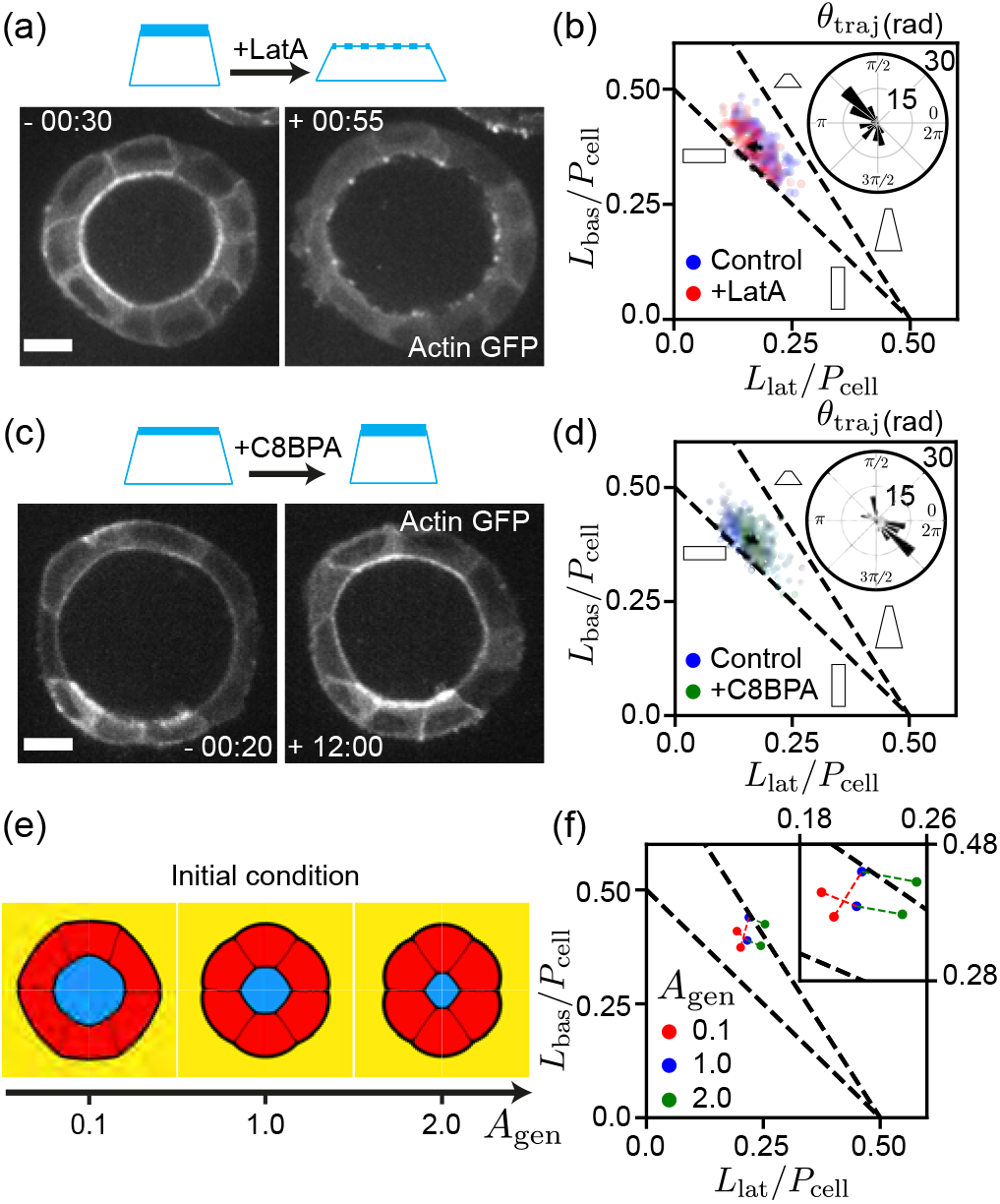
Specific modifications of actin polymerisation lead to shapes predicted by the active fluid numerical simulations. (a) Inhibition of actin polymerisation by incubating MDCK cysts with LatA. Snapshots of an individual cell before and after treatment (Video 7). (b) Cell shape diagram showing the effect of LatA on cells: cells tend to become flatter and elongate longitudinally. Black arrow represents the motion of the center of mass of the experimental cloud points before and after LatA treatment. N = 3 experiments n = 97 cells. (c) Enhancement of actin polymerisation by incubating MDCK cysts with C8BPA. Snapshots of an individual cell before and after treatment (Video 9). (d) Cell shape diagram showing the effect of C8BPA on cells: cells tend to become more radially elongated. Black arrow represents the motion of the center of mass of the experimental cloud points before and after C8BPA treatment. N = 3 experiments n = 117 cells. Insets of (b) and (d): distribution of θ_traj_, the angle of each trajectory vector in the cell shape diagram corresponding to Lat A or C8BPA treatment respectively. (e) Snapshots of simulated cysts with asymmetric turnover rates but with various baseline polymerisation rates A_gen_ (Video 10). (f) Cell shape diagram showing the effect of enhancing or diminishing active gel polymerisation in simulated cells. Insets of (b) and (d) show that distributions go in opposite directions consistently with insets of (f). Scale bars: 10 µm.

Our model also allows us to vary other parameters than those related to acto-myosin dynamics. Hence, we decreased cell-cell adhesion (M&M, Appendix. 4(e) and Appendix Fig.9). Upon treatment, cells became radially elongated and lumen occupancy decreased and these results are also substantiated by numerical simulations (Appendix Fig.9). Additionally, we changed ECM elasticity by varying the concentration of Matrigel in the culture (Appendix. 4(f) and Appendix Fig.10). Briefly, acto-myosin distributions were conserved for different concentration of Matrigel and we could reproduce the qualitative decrease in lumen occupancy at high Matrigel concentration. However, due to the numerous side effect of modifying Matrigel concentration (change in elasticity but also in available nutrients to cells and cell-ECM adhesion sites), these results call for a cautious interpretation.

Even though simulations allow us to reproduce experimental perturbations qualitatively, an exhaustive exploration of range of parameters is currently out of reach of this study and prevents further quantitative comparisons of these experiments. Also, we could not test all configurations explored in (Fig.6) with experiments available at this point. Moreover, our model does not investigate the role of tissue growth (due to cell division), an effect that could be of importance in interpreting all experiments involving timescales comparable to MDCK cell doubling time (≃ 18h [36]): for instance actin depolymerization washout (Supp. Fig. 5). In testing the myosin inhibition experiments, we were limited by the range of parameters imposed by stability of the numerical scheme (SM [32]). These points will be the focus of future studies taking into account the complexity of the cysts. However they do not challenge the relevance of our approach.

All these results confirmed that our model is fit to reproduce qualitatively a wide range of different experimental perturbations. More specifically, our cortex model coupled to a multicellular phase-field model faithfully reproduces cell shape changes obtained when varying key acto-myosin mechano-chemical parameters like polymerisation rates and acto-myosin contractility.

## III. Discussion

Asymmetric properties of the cortex are sufficient to describe cell shapes in the 3D context of cysts and organoids. The experiments identify cortical asymmetries and quantify their mechanical properties. The model then asks whether these measured asymmetries are sufficient to explain epithelial morphology. The fact that a single mechanically consistent framework reproduces a number of experimental perturbations affecting cortex dynamics but also external parameters like cell-cell adhesion and ECM elasticity, provides evidence that the proposed mechanism captures the dominant physical ingredients governing cyst morphogenesis. This framework could serve as a generic approach to design 3D tissues with molecular strategies and mesoscopic consequences on cell and cyst shapes.

This mesoscopic approach goes also with structural simplifications which may contribute to setting the contractile nature of these cortices. For example, the apical side is composed of villi with folded structures and anchors to the membrane [1]. Orientations of actin filaments are reported to be specific at this level [1]. Our results show that our mesoscopic approach is still successful to explain cell shapes beyond molecular compositions and architecture.

Cysts also emerge in *organoids* which are three-dimensional organ-models formed *in vitro* from stem cells when cultured in the relevant environments [51]. Interestingly, differentiated epithelial cells can also mimic cell shape dynamics observed in organoids. Indeed, they often share common cortical proteins as well as adhesion complexes across systems [52, 53]. These shared points allow to probe and establish phenomena with reliable extrapolation to organs found *in vivo*. Altogether the integrated comparisons and characterisation of epithelial cysts and organoids with active gels simulations appear as a powerful route to implement and test active fluid descriptions for cell shapes determination in 3D tissues. Quantitative measurement of each cell shape and mechano-chemical characterization of cells in a cyst are lacking so far.

We report that actin is primarily present in the apical side whereas acto-myosin is assembled at the basal side. One could have thought that actin gel would be extensile and acto-myosin gel contractile, like in former studies [18, 54, 55]. This asymmetry in composition does not translate into extensile and contractile gels respectively. Rather they are both contractile and this suggests that another mechanical asymmetry is at play. This also illustrates that the molecular knowledge of the gel composition is not sufficient to explain the mechanics and shape of the cortex.

Our description integrates generic cellular properties and should be broadly applicable in other organoids and in 3D tissues. It will be interesting to interrogate the cortical properties in each case to identify conservations and differences between systems.

## ACKNOWLEDGEMENTS

We acknowledge Kana Fuji and Sakurako Tanida for supporting us to implement numerical simulations. We thank the Riveline group for help and discussions and Erwan Grandgirard and the Imaging Platform of IGBMC. We also thank Anne Grapin-Botton, Masaki Sano and Alf Honigmann and their respective teams for advices and discussions. We thank Makiko Nonomura, Jacques Prost and Karsten Kruse for fruitful discussions. L.L. and T.G. are supported by HFSP and by the University of Strasbourg, and by la Fondation pour la Recherche Médicale (T.G.). T.H. is supported by Seed Fund of Mechanobiology Institute. D.R. acknowledges the Interdisciplinary Thematic Institute IMCBio, part of the ITI 2021-2028 program of the University of Strasbourg, CNRS and Inserm, which was supported by IdEx Unistra (ANR-10-IDEX-0002), and by SFRI-STRAT’US project (ANR 20-SFRI-0012) and EUR IMCBio (ANR-17-EURE-0023) under the framework of the French Investments for the Future Program. D.R. acknowledges the Research Grant from Human Frontier Science Program (Ref.-No: RGP0050/2018).

## APPENDIX 1: ADDITIONAL DATA AND QUANTIFICATION

### Appendix 1(a) - Comparison between 2D and 3D signal distributions

For each of the cell selected for analysis in the midplane of a cyst, we chose to perform the analysis in the middle plane of this cell as depicted in the scheme of Supp. Fig. 1. We made the hypothesis that intensity signals from F-actin and MRLC were invariant by rotation around an axis of symmetry perpendicular to the apical and the basal cortices and passing through their center. We give in Video 1 a visual 3D representation of the distribution of F-actin, E-cadherin and MRLC signals for a cell. In Appendix Fig.1 we show quantifications of 3D F-actin and MRLC intensity values as a function of 2D mean values for the same quantities in the midplane. We also report deviations from the midplane values and we show that choosing the averaged midplane intensity value as a proxy for the intensity values in each cell is a reasonable hypothesis even though a systematic bias can be measured between 2D and 3D measurements.

**Appendix FIG. 1.**
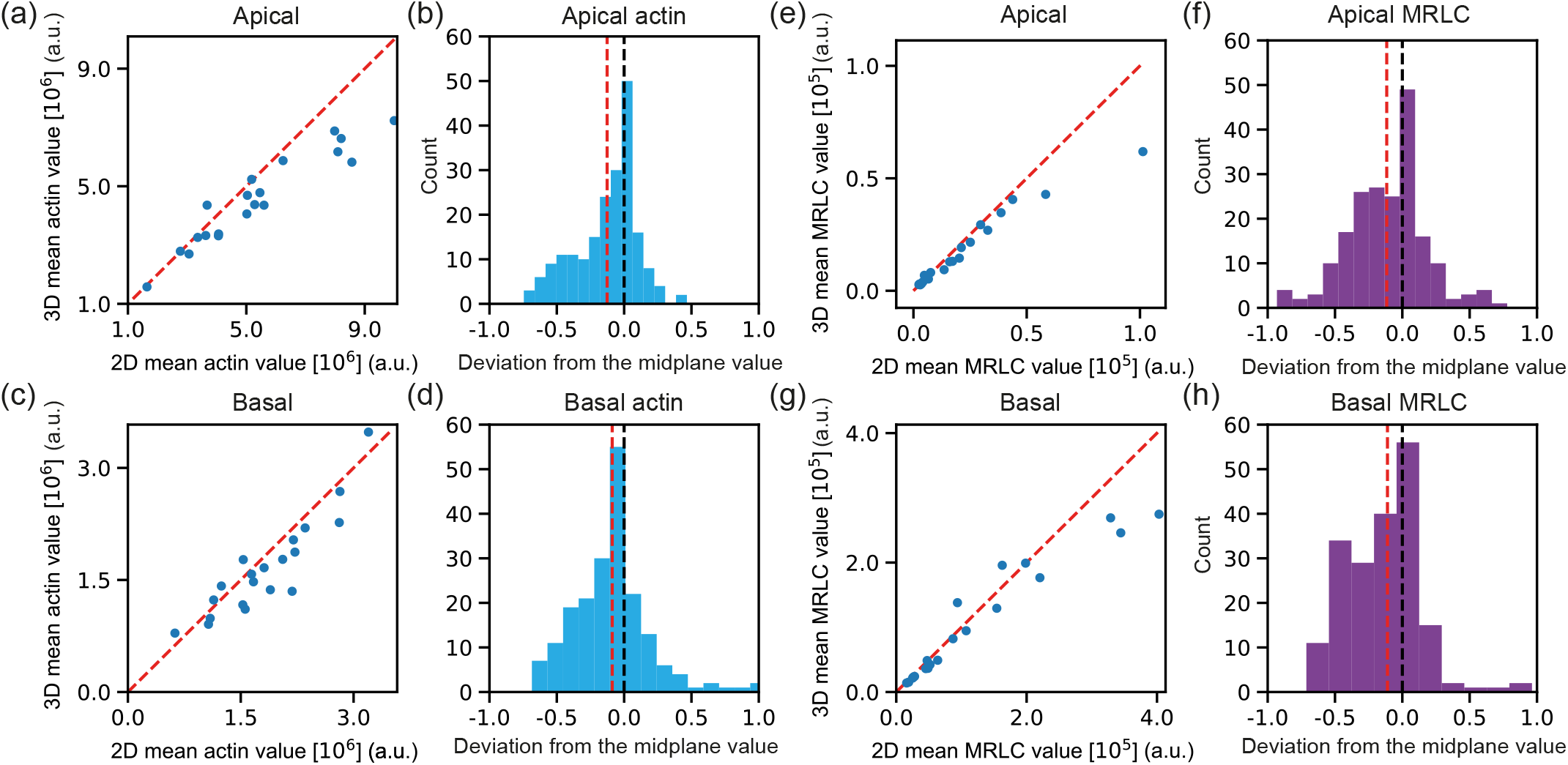
Comparison between 3D mean values and 2D intersection. - (a,c,e,g) Signal intensity value averaged along several sections as a function of signal intensity value in the midplane of the cell. (a) apical actin, (c) basal actin, (e) apical myosin, (g) basal myosin. (b,d,f,h) Histograms showing distributions across the z-axis. The horizontal axis is normalized by the 2D mean value, i.e., mean signal intensity value in the midplane. (a.u.) - red dashed line : mean of the distribution - black dashed line 0.0 value. (b) apical actin, (d) basal actin, (f) apical myosin, (h) basal myosin.

We performed complementary analysis and plotted the distribution of ratio of 2D over 3D measurements (Supp. Fig. 2 (a-g)). For apical/basal F-actin/MRLC, we systematically detected a bias indicating an overestimation of 2D measurements in the midplane of the cell over the 3D average values of ~10 − 15% (Supp. Fig. 2 (h-k)). A potential explanation for this bias can be the exponential loss of signal associated to scattering of light along the z-axis away from the objective resulting in a lower averaged value when averaging values above (with lower signal than the middle plane) and below (with higher signal than the middle plane not compensating the loss of signal in the other part).

To further evaluate if this difference is dependent on the intensity of the measured signals, we used a Bland-Altman test [56] designed for this type of evaluation. We find a significant dependence on signal intensity for apical F-actin and apical MRLC measurements (with respectively a slope of 0.2 and 0.1 respectively) but not for basal F-actin and basal MRLC (Supp. Fig. 2 (h-k)).

The dependence for apical actin and MRLC could be related to the way F-actin is organized within the apical cortex. At this location, F-actin is stored in microvilli which are densely packed [57]. A middle cut captures them in the optimal configuration where they are all aligned, thus increasing the measured signal value.

### Appendix 1(b) - Cortical width and curvature

Appendix Fig.2(d) shows the measurements of apical, basal and lateral cortices width distributions. These measurements confirmed that the apical cortex was thicker than the basal one which was itself thicker that the lateral cortex.

Appendix Fig.2(e) shows the measurements of apical, basal and lateral curvatures. We see that while the basal cortices are always positively curved towards the interior of the cyst, the apical cortices can be found to be positively or negatively correlated.

Interestingly, the lateral cortices were measured to be in some cases more curved than the apical or basal cortices. This is mainly due to the presence of highly curved region on lateral cortices mainly at the junction with the basal cortices (see E-cadherin staining in Appendix Fig.3-day7.

### Appendix 1(c) - Myosin and actin fluorescence intensity quantifications

Appendix Fig.2(b-c) show the measurements of apical to basal intensity ratio for F-actin (cyan) and MRLC (magenta). For measurements of mean values of intensity ratios, refer to Table I.

### Appendix 1(d) - Correlations between cell shape and other morphological quantities

#### Lumen occupancy

As presented in Fig.3(c-d) of the main text, we used a cell shape diagram to represent the measurements performed on single 2D cuts of cells in 5-day-old MDCK cysts. Appendix Fig.6(a) shows the correlation between lumen occupancy, *i.e* the ratio between lumen volume and cyst volume, and cell shape: *L*_occ_. Since we measure quantities in 2D, we measure lumen occupancy from surfaces. As exemplified in Appendix Fig.6(a), the larger the lumen occupancy, the flatter and the less columnar the cell.

Appendix Fig.6(b) shows the correlation between lumen occupancy and cell shape for a theoretical ideal case of a spheroid of radius *R*_*s*_, whose middle plane is composed of *N* cells surrounding a lumen of radius *R*_*ℓ*_. In this case we have simply: 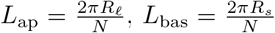 and *L*_lat_ = *R*_*s*_ − *R*_*ℓ*_. We represent the cases for *N* = 8 and *N* = 16 cells Appendix Fig.6(b) with good qualitative comparison with panel (a). Additionnaly, we have quantified how distributed the cell shape parameter *α* is w.r.t its geometrical counterpart *α*_geometrical_ = *R*_*ℓ*_*/R*_*s*_, defined by assuming a unique cell shape within a cyst (Supp. Fig. 3). This graph shows that *α* is adapted to fairly capture the heterogeneity in cell shape.

**Appendix FIG. 2.**
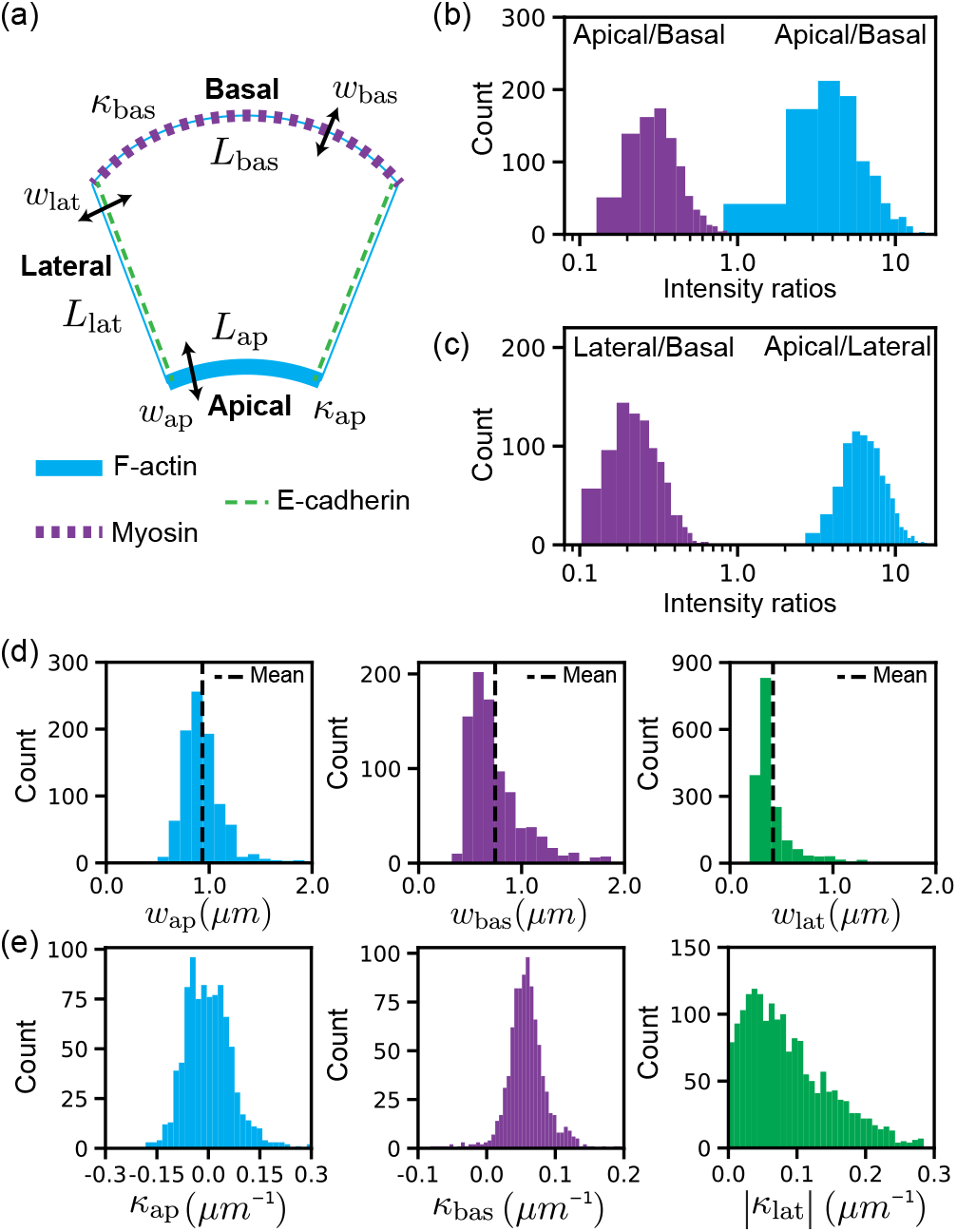
Additional quantifications of actin and myosin distributions and cell shape descriptors. - (a) Typical mean cell description that is used to quantify the relevant geometrical quantities to describe the cell cortex and the cell shape. Quantification of (b) apical to basal intensity ratios, (c) lateral to basal and apical to lateral intensity ratios for F-actin (blue) and myosin (magenta) intensity signals. (d) Quantification of apical, basal and lateral acto-myosin cortex widths. (e) Quantification of apical, basal and lateral acto-myosin cortex curvatures. N = 3 experiments, n = 791 cells.

#### Curvatures

Appendix Fig.6(c-d) represent the correlation of apical and basal curvatures with the cell shape. Positively curved apical lines correspond to elongated cells (high *L*_lat_*/P*_cell_ ratios) when basal curvatures show no strong correlations with cell shape.

### Appendix 1(d) - Conservation of acto-myosin asymmetry distributions and correlation with cell shape

Appendix Fig.3 represents MDCK cysts fixed and stained at day 3, 5 and 7 days of culture for E-cadherin, F-actin and MRLC. This panel shows that the asymmetry of F-actin and MRLC distributions is conserved over time. The magnified box on individual cells of the different cysts represent typical examples of cells in these experimental conditions. Quantifications of correlations in the cell shape diagram for day 3, 5 and 7 as presented in Fig.3 are shown in Appendix Fig.4. Note that the lumen size gradually increases between day 3 and day 7 due to water transport by osmotic pressure inbalance [58].

In Appendix Fig.5 we plot the distance between each point in the cell shape diagrams and the line defined for 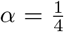 as a function of intensity ratios for either F-actin or MRLC. To test for monotonic increase or decrease, we computed Spearman’s rank correlations for all conditions.

## APPENDIX 2 QUANTIFICATION OF MECHANO-CHEMICAL PROPERTIES OF THE ACTO-MYOSIN CELL CORTEX

### Appendix 2(a) - FRAP experimental setup

We controlled the microscope using the FRAP module on the Metamorph software (iLas2). A calibration step was needed to align the FRAP laser with the objective, for this purpose, we used a glass bottom Petri Dish on which we draw a line of fluorescent marker that then served as target to calibrate the FRAP module. This step was crucial for getting reproducible laser shots from the system. We defined a circular (2*µ*m diameter) ROI to be photobleached using the iLas2 software. We repeatedly photobleached a ROI 50-100 times for a total exposure time of 185-370 ms with a laser (488 nm) at 15% of its nominal power. We also took care of placing the analysed cysts at the center of the field of view. We recorded the dynamics of recovery every 0.5s for 75s and then every 5s for 200s.

**Appendix FIG. 3.**
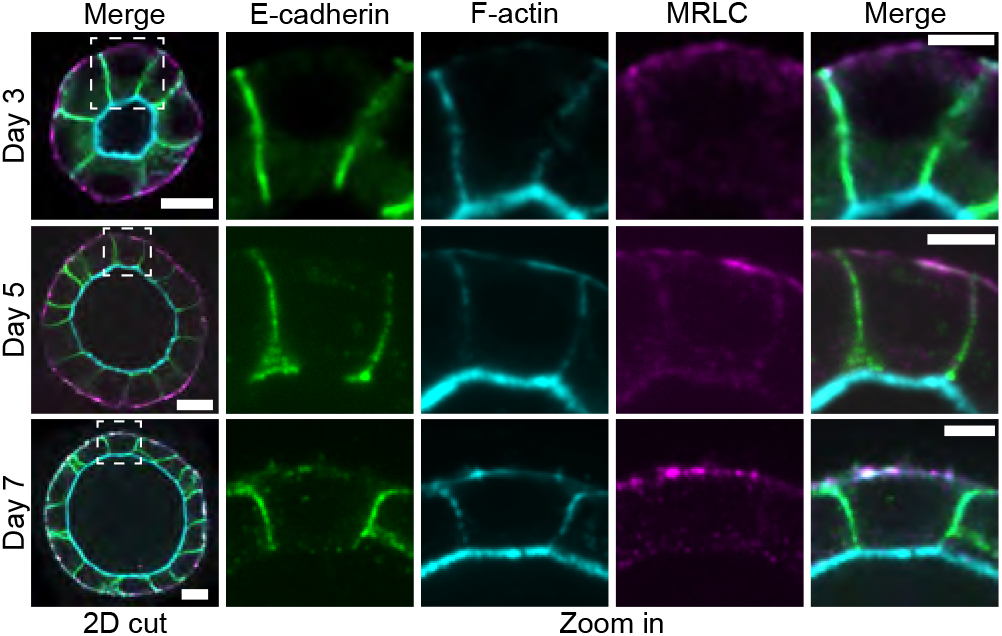
Distributions of actin and myosin in MDCK cysts are conserved over **time**. - (a) 3-, 5- and 7-day-old MDCK spheroids embedded in Matrigel, fixed and stained for F-actin (phalloidin, cyan), MRLC (magenta) and E-cadherin (green). We also provide magnified images of the areas highlighted by a box. The apical side of each spheroid exhibits a thick layer of F-actin and less actin on the lateral sides and basal sides. Myosin distribution shows an increase in clusters at the basal side. Scalebars: 10 µm in cyst snapshots, 5 µm in cells zoom in.

### Appendix 2(b) - FRAP quantification

To analyse FRAP experiments, we measured the average signal in the bleached ROI over time (Fig.4(b)), in a background region and in a ROI in the cytoplasm using a FIJI custom code. We subtracted the mean intensity of the background taken outside of the cyst to the intensity in the ROI overtime *I*_FRAP_ = *I*_ROI_ − *I*_back_. Then we renormalised the signal in order to compare experiments between each other by computing:

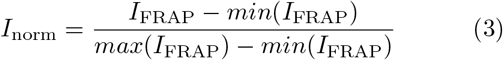

**Appendix FIG. 4.**
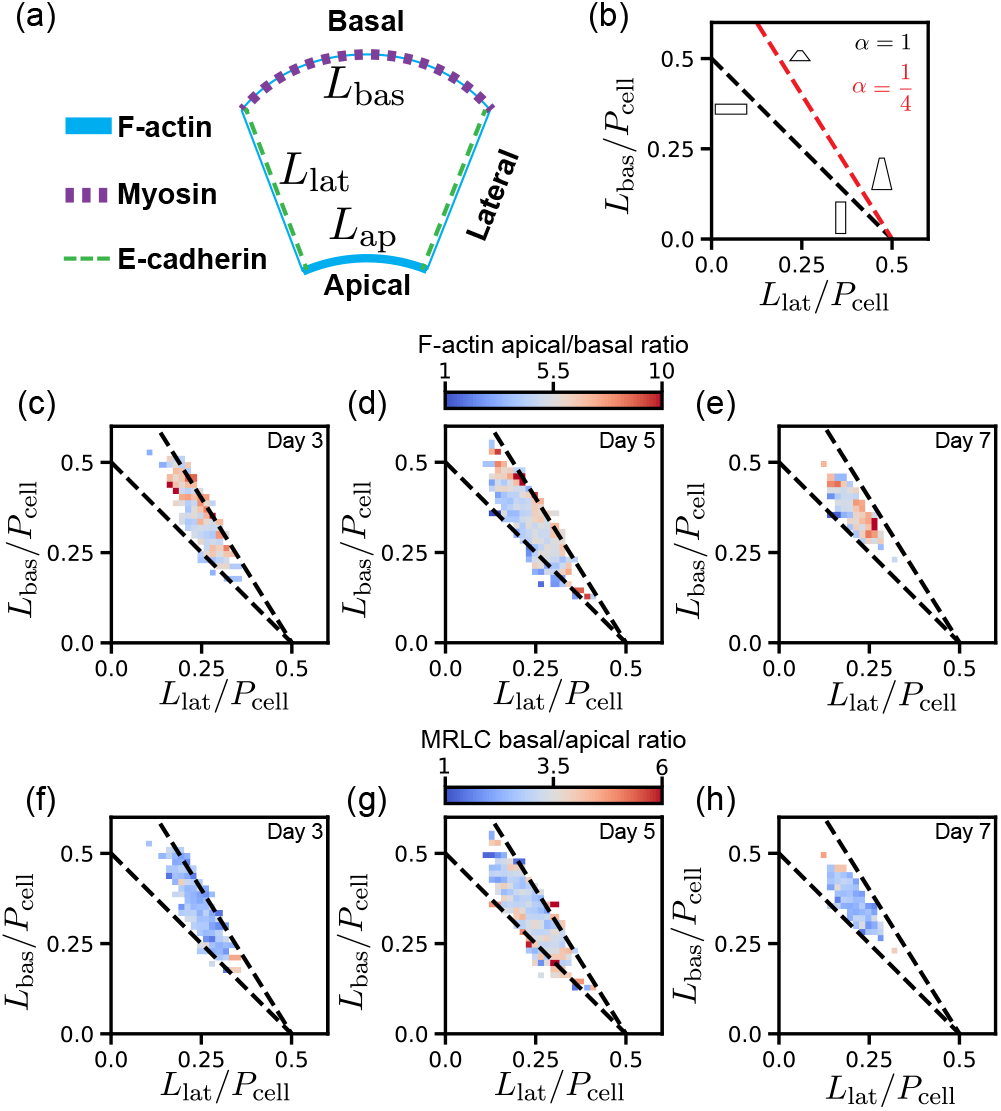
Correlation between acto-myosin cortical density and cell shape in MDCK cysts. - (a) Typical mean cell description used to quantify the relevant geometrical quantities to describe the cell cortex and the cell shape. (b) Schematic description of the cell shape diagram. Oblique lines correspond to different apical to basal length ratios α as described in the main text. Moving along the line changes the aspect ratio of cells w.r.t. the lateral length. F-actin apical to basal intensity ratio correlation with cell shape at day 3 (c), day 5 (d) and day 7 (e). MRLC basal to apical intensity ratio correlation with cell shape at day 3 (f), day 5 (g) and day 7 (h). P_cell_, cell perimeter. For all measurements, N = 3 experiments, n = 258 cells quantified in 3-day-old MDCK cysts, n = 905 cells quantified in 5-day-old MDCK cysts, n = 158 cells quantified in 7-day-old MDCK cysts.

Next we fitted the curve using a non linear least square method for a function 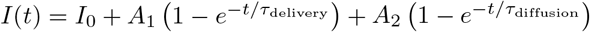 (Fig.4(c-f)) and [59]). From the fit, we obtained the values of (*I*_0_, *A*_1_, *A*_2_, *τ*_delivery_, *τ*_diffusion_). By definition, the mobile fraction is: *M* = 1™(*I*_0_ +*A*_1_ + *A*_2_).

### Appendix 2(c) - FRAP additional analys

#### Mobile fracti

The mobile fraction quantifies the portion of molecules that is renewed after recovery fol lowing the photobleaching. We measured that the mobile fraction of actin molecules in the apical cortex was lower than in the basal and lateral cortices which showed similar values for this measurement. This result can also be seen by comparing Fig.4(d-f) where the recovery dynamics for apical, basal and lateral actin normalised intensity are shown. This result tends to indicate that the geometry of the actin cortex at the apical side makes it more difficult for G-actin molecules to be replaced after photo bleaching. This is consistent with the dense actin region reported in villi at the apical cortex of MDCK cells in cysts [60]

#### Diffusion time

We estimated the diffusion times for the apical, lateral and basal cortices. We found similar times of the order of a few seconds (Fig.4(g)), which correspond to the typical diffusion time of G-actin monomer, as was previously measured in [48, 49, 60, 61].

**Appendix FIG. 5.**
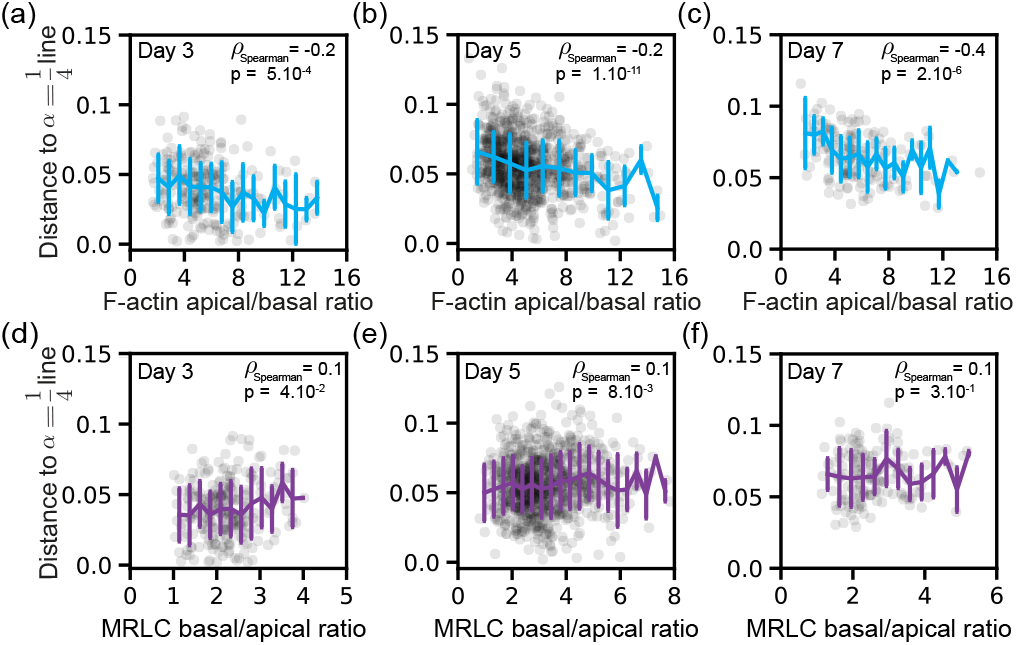
Additional quantifications on the correlation between acto-myosin cortical density and cell shape in MDCK cysts. - Distance to the alpha 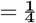 line in the cell shape diagram as a function of F-actin apical/basal intensity ratio (a, b, c) or MRLC basal/apical intensity ratios (d, e, f) for day 3, day 5 and day 7 respectively. Spearman’s rank correlation tests with their p-values are indicated in the top right corner of each plot. For all measurements, N = 3 experiments, n = 258 cells quantified in 3-day-old MDCK cysts, n = 905 cells quantified in 5-day-old MDCK cysts, n = 158 cells quantified in 7-day-old MDCK cysts.

**Appendix FIG. 6.**
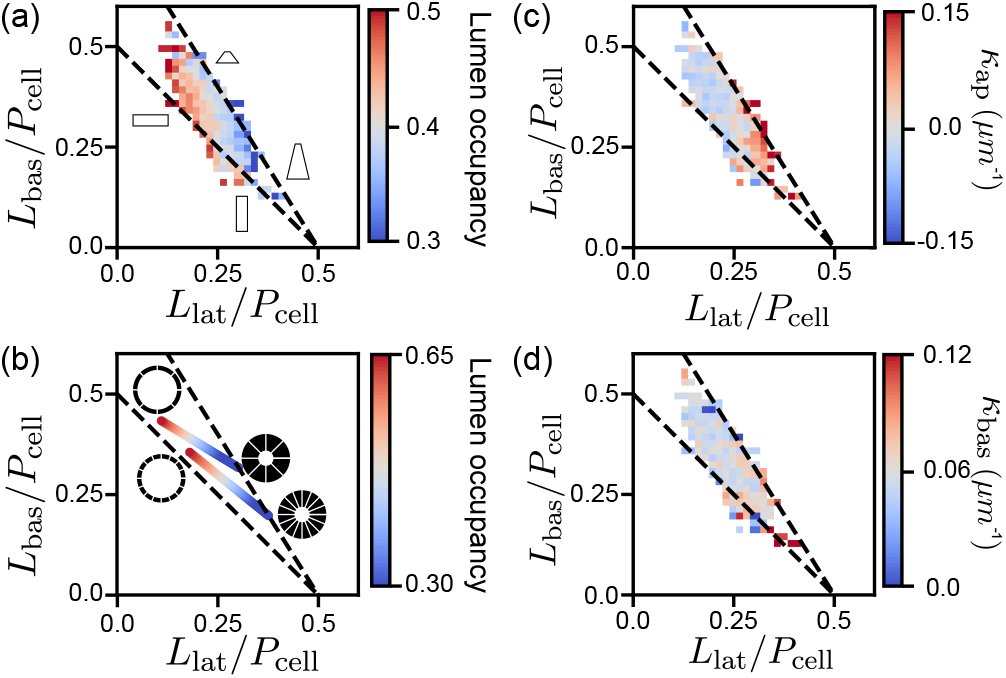
Correlation between cell shape, lumen occupancy and apical and basal cortex curvatures in MDCK cysts. - (a) Correlation of lumen occupancy with cell shape. (b) Correlation of lumen occupancy with cell shape - Theoretical results obtained for a symmetric configuration of a cyst containing 8 cells (top line) or 16 cells (bottom line). Apical curvature correlation with cell shape. (d) Basal curvature correlation with cell shape. For all measurements, N = 3 experiments, (a-c-d-e(middle)) n = 791 cells; (e(left)) n = 258 cells; (e(right)) n = 158 cells.

### Appendix 2(d) - Laser ablation experimental setup

A typical radius of a cyst at Day 5 is about 0.05 mm, so using the double layer Matrigel protocol was key in performing these laser ablation experiments and not be bothered by the working distance of the objective. By adjusting the scanning window size to 512 x 120 pixels, it was possible to reduce the frame interval to a minimal value of 0.24 s (Fig.5) (or to 256 x 120 pixels - 0.12s frame interval in Supp. Fig. 4) between frames to follow the ablated regions of the cortex. The laser is a Ti:Saph pulsed laser (Coherent Chameleon Ultra II) used at the 800nm wavelength through an objective Leica Plan Apochromat 63X oil 1.4 NA. It emits ~100fs, at the repetition rate of 80MHz, mean power of the laser ~4W. For the parameters used for ablation, it corresponds to a mean power of the laser ranging between 80 mW and 300 mW measured at the back focal plane of the objective. For detection of the signal, we used a Hybrid Detector (HyD) descanned detector for which we adjusted the gain to maximise the signal to noise ratio for each ablated regions. After characterisation of the MP laser and tests, the offset value was set to 35%. Then, depending on the depth of the sample to ablate, parameters for proper ablation were found through trial-and-error. Typical working parameters for ablation of the basal cortex of MDCK cells in cysts are: transmission T = 50% and gain G = 45%.

### Appendix 2(e) - Laser ablation additional analysis

We detail here the additional results shown for the laser ablation experiments and their interpretation in term of inference of cortical stresses. We performed laser ablation experiments [26] to test the contractile or extensile nature of the active stresses generated by actin and myosin within the different parts of the cell cortex. When ablating apical, basal or lateral cortices, we observed an opening of the actin cortex of a few micrometers within seconds (Fig.5(b-d)) and Supp. Fig. 4). These openings were larger in the case of the basal cortex (Video 4, Fig.5(c,e)) than in the case of the apical (Video 3, Fig.5(b,e)) or the lateral (Video 5, Fig.5(d,e)) cortices. This demonstrates that the cortices are more tensile at the basal sides rather than at the apical and lateral sides. We quantified the opening velocity 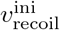 after laser ablation as a readout for comparing stress between the different apical, lateral and basal cortices, assuming they have similar damping coefficients [26]. Even though these laser ablation experiments present limitations (see [62, 63]), they represent the most reliable and common approach at this stage.

**Appendix FIG. 7.**
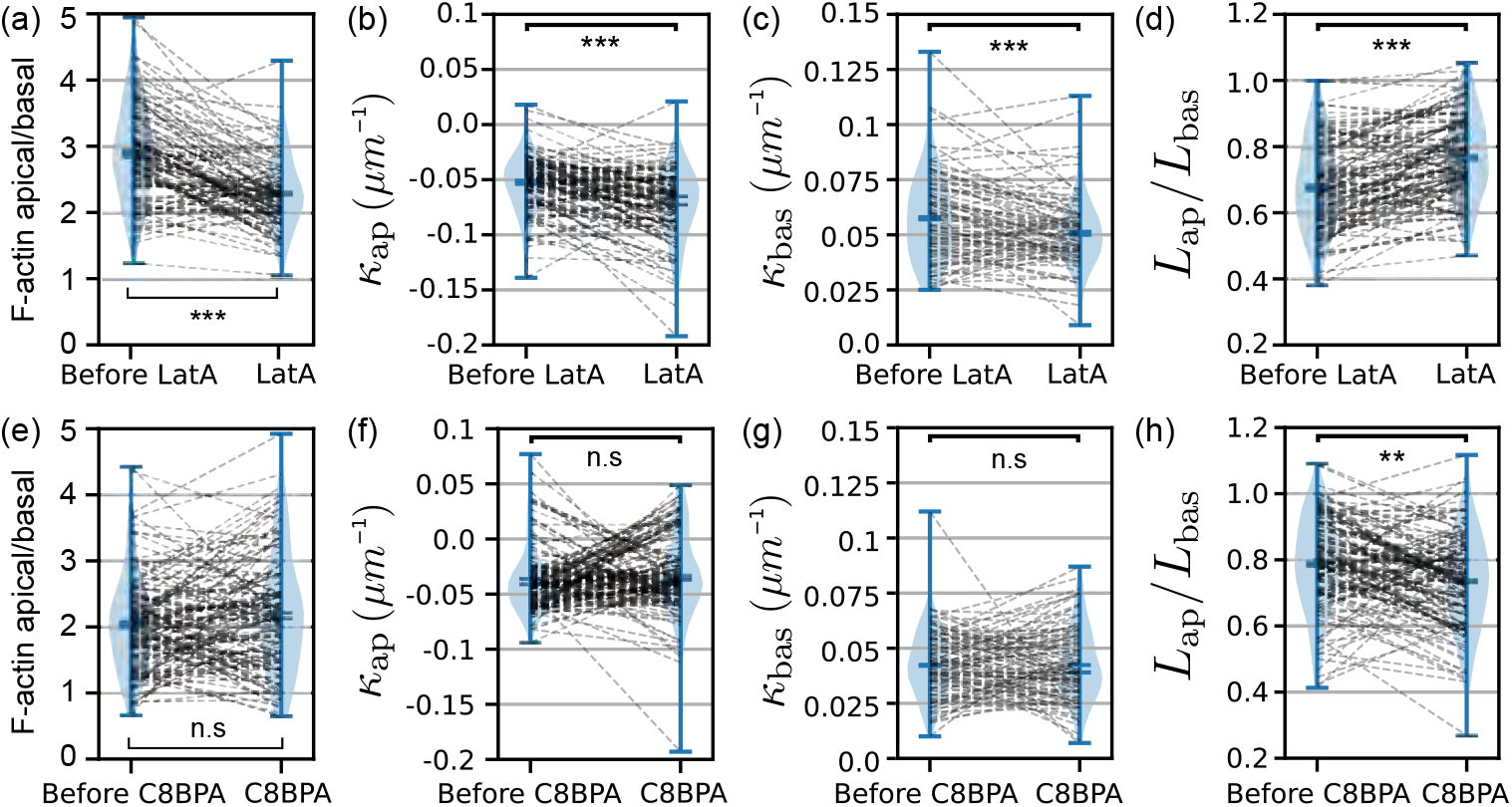
Additional quantifications following drug treatments with Latrunculin A and C8BPA. - Quantification of (a) changes in apical to basal actin intensity ratios, (b) apical curvature, (c) basal curvature, (d) ratio of apical length to basal length, before and after treatment with 200 nM of Lat A. N = 3 experiments, n = 97 cells. Quantification of (e) changes in apical to basal actin intensity ratios, (f) apical curvature, (g) basal curvature, (h) ratio of apical to basal lengths before and after the treatments C8BPA: N = 3 experiments, n = 117 cells. Statistical tests: *:p < 0.05, **: p < 0.01, ***: p < 0.001, n.s: p > 0.05.

## APPENDIX 3 CORRESPONDANCE BETWEEN EXPERIMENTAL AND NUMERICAL PARAMETERS

For Fig.6(g), we chose parameters fitting the experimental characterisation that we performed in Fig.4, Fig.5 and Table I.

From the FRAP experiments, we measured (Fig.4 and Table I):

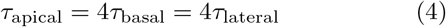

Because we probed the dynamics of assembly and dissassembly of the actin network with FRAP experiments, we translated these measurements into values of *A*_deg_:

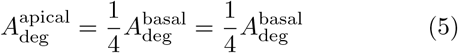

From the actin intensity measurements we measured (Fig.3, Appendix Fig.2 and Table I) assuming that intensities of fluorescently labelled actin is a readout for actin density:

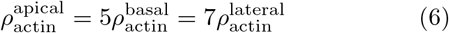

To use this information, we made the hypothesis that the measurements reflected the steady-state density of the actin cortex. We could then write the conservation equation for the active gel density (Eq.(18) in the SM[32]) at steady state (with no actin flows) and find that:

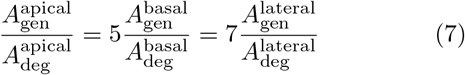

**Appendix FIG. 8.**
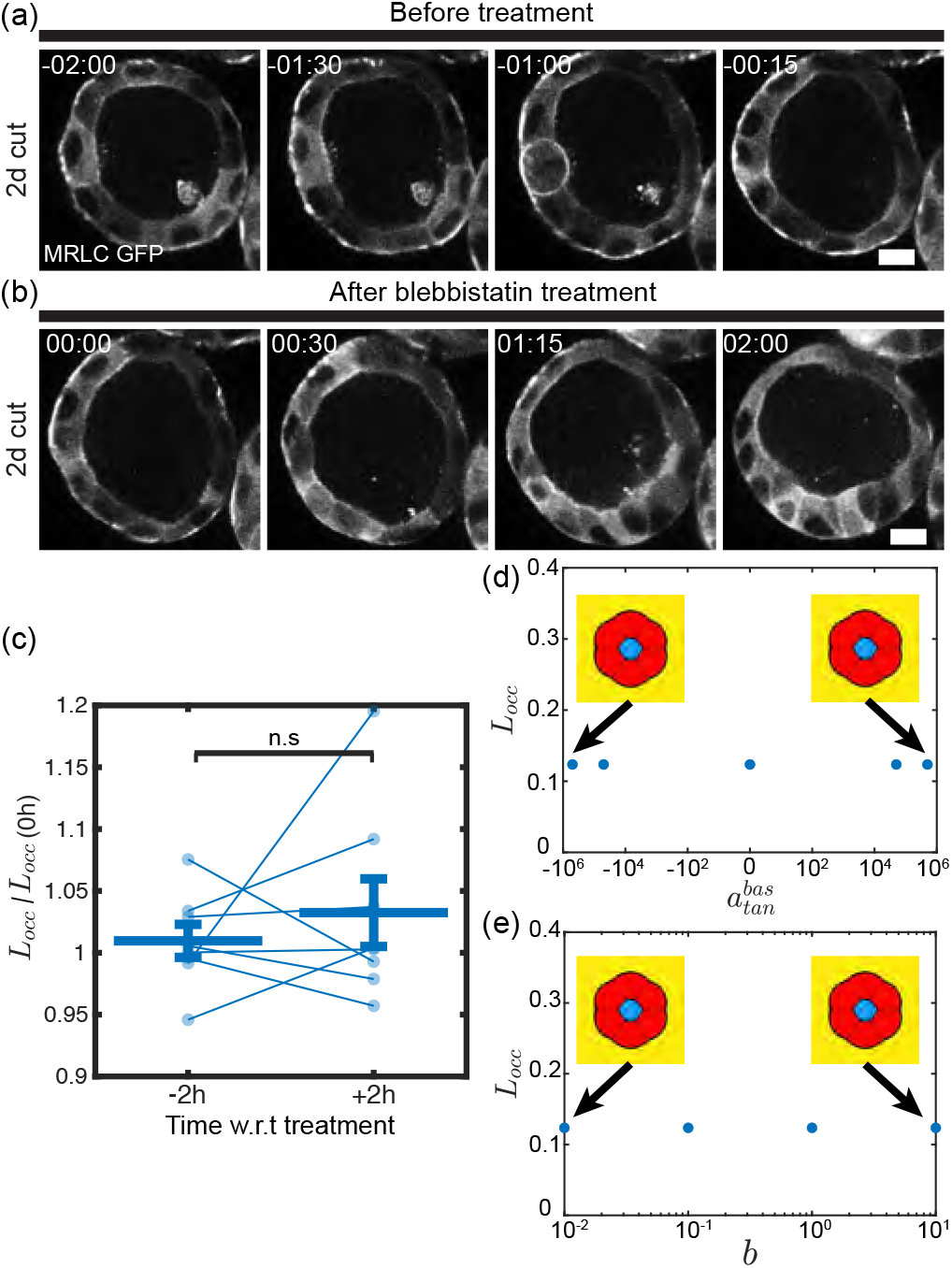
Inhibition in myosin activity does not change cell shapes in agreement to simulation. To investigate the role of myosin in regulating cell shape and cyst architecture, day 5 cysts were treated with the myosin II inhibitor blebbistatin (c = 10 µM final concentration). (a-b) Representative snapshots show a cyst before (a) and after (b) blebbistatin treatment, visualized with MRLC-GFP (grey). Time in hh:mm, scale bar: 10 µm. (c) Measurement of variations in lumen occupancy 2h before and 2h after blebbistatin treatment. (d-e) Sensitivity parameter analysis for parameters related to active gel contractility (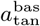 and b) shows that, in the given range of tested parameters, a variation in these parameters led to no changes in lumen occupancy (panels (d) and (e)).

By combining these two results we got:

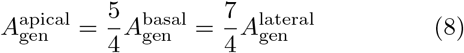

The laser ablation experiments provided insight into the cortical tension at apical, lateral and basal sides of the cells (Fig.5 and Table I). These measurements can serve as estimate for the active contractile stresses. The experiments provided ratios of contractile tensions through measurements of initial recoil velocity [26]:

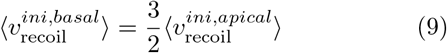

To estimate ratios of contractile stresses we made the hypothesis that the contractile stresses measured by laser ablation represented the tangential active stresses in the active gel model. Then we had from the definition of Π(*ρ*):

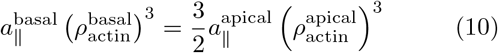

Using the previous measured actin densities, we estimated that:

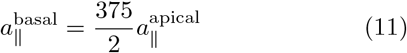

For the exact choice of parameters, refer to SM [32].

## APPENDIX 4 PERTURBATION EXPERIMENTS

We detail here the additional quantifications of the drug experiments targetting actin polymerization, myosin contractility and cell-cell adhesion. For the respective protocols, see M&M.

### Appendix 4(a): Quantification of cell shape changes

To quantify cell shape changes, we used the cell shape diagram 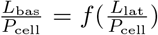 defined in Fig.3(b). We defined *θ* = arctan(*y*_2_ − *y*_1_, *x*_2_ − *x*_1_) as the angle of a trajectory vector connecting two points (M_1_(*x*_1_, *y*_1_) and M_2_(*x*_2_, *y*_2_)) in this diagram and arctan(x, y) is the arc tangent of y/x (taking into account in which quadrant the point is located).

This angle allows to visualise how drug treatments modify cell shape and in which ‘direction’ a cell evolves in this diagram following a drug treatment (insets of Fig.7(b,d) and Supp. Fig. 5).

### Appendix 4(b): Depolymerizing F-actin

We incubated 5-day-old MDCK cysts with a depolymerizing actin drug LatA at a concentration of *c* = 200 nM and we monitored the change in cell shape up until 1 hour of incubation (Video 7). After 1h, we washed out LatA and changed back to fresh culture medium (M&M and Video 8). We quantified the changes in apical and basal curvatures (Appendix Fig.7(b-c)) as well as the ratio between apical and basal length (Appendix Fig.7(d)). Both apical and basal curvatures, as well as apical to basal length ratio increased in absolute values indicating that cells got deformed by a radial pushing force (*i.e* lumen hydrostatic pressure) following the depolymerisation of the actin cytoskleton upon LatA treatment.

**Appendix FIG. 9.**
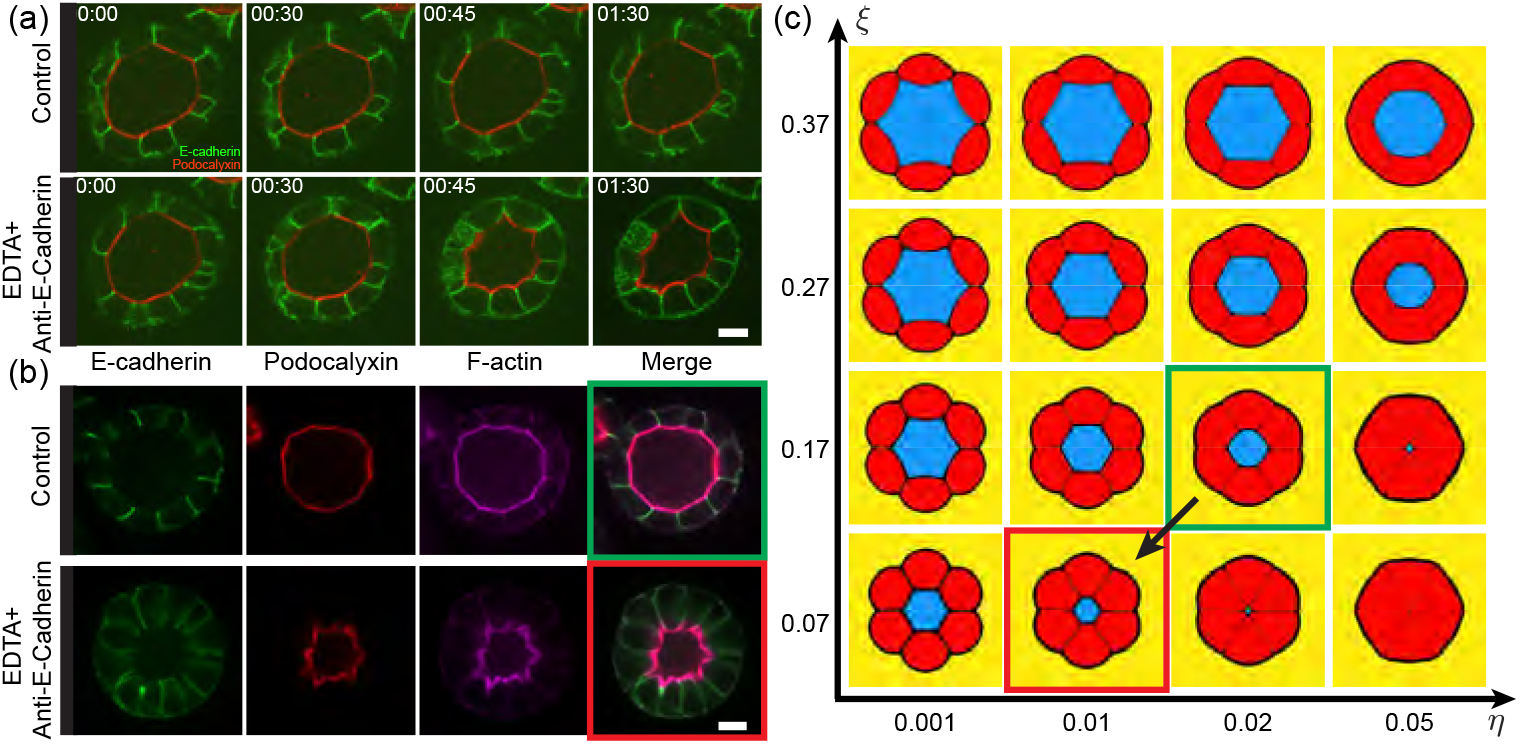
Decrease in cell-cell adhesion changes cell phenotype. (a) Cysts were incubated with function-blocking antibodies against E-cadherin in the presence of EDTA to disrupt calcium-dependent adhesion [36–38]. Time-lapse snapshots show control cysts (top) and cysts treated with EDTA and E-cadherin antibodies (bottom). E-cadherin is shown ein green; podocalyxin in red. (b) Representative snapshots of fixed cysts showing the distribution of E-cadherin (green), podocalyxin (red), and F-actin (magenta), along with merged channels. Control (top) and EDTA + antibody-treated (bottom) conditions are shown. Disruption of E-cadherin-mediated adhesion leads to distinct changes in cyst cell morphology compared to controls. scale bar: 10 µm (c) Phase diagram over lumen pressure and cell-cell adhesion predicted by the simulations. The changes in phenotype observed in (a-b) were matched to their guessed counterpart in simulations. Starting configuration : (η = 0.02, ξ = 0.17) is outlined in green and the final phenotype is outlined in red. Black arrow represents the evolution of parameters reproducing the experimental perturbation of lowering cell-cell adhesion. Of note: cell-cell adhesion η may be decreased together with osmotic pressure ξ, because the opening of junctions affects also osmotic pressure [36, 65].

In Supp. Fig. 5, we show and quantify the change in cell shape following LatA washout (quantifications performed after 11 h of washout for a typical experiment). The washout of LatA induced a change in cell shape from an elongated cell to a more cuboidal one (Supp. Fig. 5 (b)). In this case, the actin ratio between the apical and basal cortices (Supp. Fig. 5 (c)), the ratio between the apical and basal cortical lengths (Supp. Fig. 5 (d)) and apical and basal curvatures (Supp. Fig. 5 (e-f)) came back to similar levels as prior LatA treatment.

Altogether, the washing out of LatA induced a polymerization of actin filaments in the acto-myosin cortices and a transition back to the initial cellular shape within MDCK spheroids.

As discussed in [64], lumen occupancy of the cyst and the volume of the luminal cavity seems to be closely linked to the actin polymerisation in the cellular cortex.

### Appendix 4(c): Enhancing F-actin polymerization

We incubated 10-day-old MDCK cysts with *c* = 100 *µ*M of an enhancer of actin polymerization C8BPA (M&M) and monitored the change in cell shape up until 12 hours of incubation (Video 9). We quantified the changes in apical and basal curvatures (Appendix Fig.7(fg)) as well as the ratio between apical and basal length (Appendix Fig.7(h)). Even though apical and basal curvatures did not evolve on average, apical to basal length ratio decreased indicating that an increase in actin polymerisation resulted in cells getting more elongated and cuboidal.

### Appendix 4(d): Inhibiting myosin

We incubated 5-day-old MDCK cysts with *c* = 10 *µ*M of myosin inhibitor Blebbistatin (M&M) and monitored the change in lumen occupancy between 2h hours before treatment and 2 hours after treatment (Appendix Fig.8(h)). We also provide a sensitivity analysis for the parameters related to the contractility of the active gel in our model (Supp. Fig. 6, Supp. Fig. 7 and Supp. Table 2). Inhibition of myosin does not affect cell shape and cyst geometry in experiments and in simulations.

### Appendix 4(e): Decreasing cell-cell adhesion

To decrease cadherin-mediated adhesion we used 5 mM EDTA and an anti E-cadherin blocking antibody at a concentration of 10 *µ*g/ml (M&M). Upon treatment, cells became radially elongated and lumen occupancy decreased (Appendix Fig.9(h)(a,b)). These results were compared with numerical simulations in which we varied cell-cell adhesion *η* as well as lumen pressure *ξ*. Interestingly, decreasing cell-cell adhesion in MDCK cysts not only change the strength of cell-cell contact but also increases the fluid leakage in the paracellular region leading to a decrease in lumen volume. We recapitulate this result in the numerical phase diagram (Appendix Fig.9(h)(c)).

**Appendix FIG. 10.**
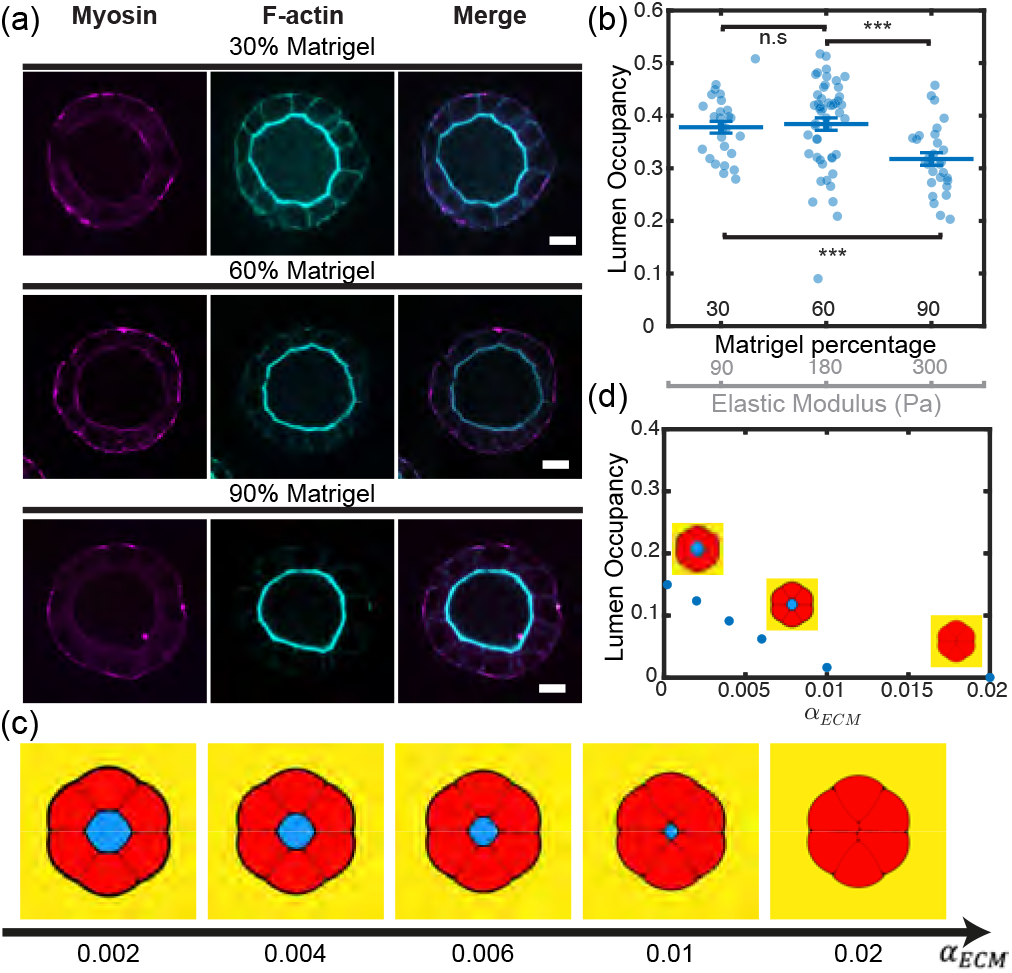
Effect of Matrigel concentration on cell and cyst phenotypes. (a) Cysts were cultured in Matrigel at varying concentrations: 30% (top row), 60% (middle row), and 90% (bottom row). For each condition, images are shown from left to right as follows: myosin (magenta), F-actin (cyan), and merged channels. Scale bar: 10 µm. (b) Lumen occupancy as a function of Matrigel percentage. Matrigel percentage are translated into Young Modulus by using [66]. (c) Starting from a steady-state, effect of varying ECM elasticity on simulated cyst phenotypes (snapshot at 100000 numerical integration steps). (d) Lumen occupancy as a function of ECM elasticity in simulations. Distributions of actin and myosin were similar but the trend in the lumen occupancy was changed which is consistent with numerical simulations.

### Appendix 4(f): Varying Matrigel concentration, ECM elastic modulus

We changed the culture condition of MDCK cysts by using diluted Matrigel of different concentrations. Of note, this method does not guarantee to have an homogeneous ECM around each cysts since Matrigel will sink by gravity in the culture dish.

The asymmetry in actin and myosin distribution is conserved for all conditions (Appendix Fig.10(a)). In addition, increasing the ECM elasticity parameter in our simulation led to a decrease in lumen occupancy and radial cell elongation, consistent with experiments (Appendix Fig.10(c)). In contrast to simulation, in experiments a change in Matrigel may have multiple effects, i.e. changes in elasticity but also changes in nutrients as well as density of anchoring points. This makes this experiment difficult to use as a test in our context.

To estimate the change in elastic modulus, we refer to the calibration curve relating Matrigel concentration to its elastic modulus available from the producing company Corning [66].

