## Supplementary Material for "Asymmetric cortical mechanics shape epithelial cell geometry in organoids"

### 1 Supplementary materials

#### 1.1 Code availability

The code to simulate the multicellular phase field coupled to an active gel cortex can be found at `https://github.com/tristanguyomar/multicellular_phase_field_active_gel_organoids`.

#### 1.2 Supplementary videos

- **Supplementary Video 1 - 3D rendering of MDCK cell in a 5-day-old cyst**: 3D rendering of a single MDCK cell isolated from a 5-day-old MDCK cyst embedded in Matrigel. E-cadherin (green), F-actin (cyan), and myosin MRLC (magenta). Top: X-axis rotation view, bottom: Y-axis rotation. Scalebar: 5 $\mu$m.
- **Supplementary Video 2 - FRAP experiments to probe actin dynamics in apical basal and lateral cortices**: Representative examples of FRAP experiments performed on apical, basal and lateral cortices. MDCK cells expressing actin-GFP in 5-day-old cysts. Scalebar: 5 $\mu$m. Time in mm:ss.
- **Supplementary Video 3 - Laser ablation of apical cortices**: Representative examples of laser ablation of the apical cortex. MDCK cells expressing actin-GFP in 5-day-old cysts. Time in ms. Scalebar: 5 $\mu$m.
- **Supplementary Video 4 - Laser ablation of basal cortices**: Representative examples of laser ablation of the basal cortex. MDCK cells expressing actin-GFP in 5-day-old cysts. Time in ms. Scalebar: 5 $\mu$m.
- **Supplementary Video 5 - Laser ablation of lateral cortices**: Representative examples of laser ablation of the lateral cortex. MDCK cells expressing actin-GFP in 5-day-old cysts. Time in ms. Scalebar: 5 $\mu$m.
- **Supplementary Video 6 - Representative movie of a multicellular phase field coupled to active gel cortices simulation**: Timelapse of a simulated spheroid initiated from 6 cells. Parameters used to generate this movie are listed in subsection 4.3.5. Left: ECM phase-field (yellow), cells phase-fields (each in red), lumen phase field (blue). Right: Active gel density map.
- **Supplementary Video 7 - Latrunculin A treatment**: MDCK cells expressing actin-GFP in 5-day-old cysts treated with 200 nM of Latrunculin A. Time in mm:ss. Scalebar: 10 $\mu$m.
- **Supplementary Video 8 - Latrunculin A treatment and washout of 5-day-old cysts**: MDCK cells expressing actin-GFP in 5-day-old cysts treated with 200 nM of Latrunculin A followed by washout. Time in hh:mm. Scalebar: 10 $\mu$m.

- **Supplementary Video 9 - C8BPA treatment of 10-day-old MDCK cysts**: MDCK cells expressing actin-GFP in 10-day-old cysts treated with 100 $\mu$M of C8BPA. Time in hh:mm. Scalebar: 5 $\mu$m.

- **Supplementary Video 10 - Active gel turnover rate perturbation leads to cell and organoid morphological changes.** Movie illustrating the numerical simulations performed to generate Fig. 4(e-f): the baseline active gel polymerisation rates $A_{\text{gen}}$ are indicated below each condition. ECM phase-field (yellow), cells phase-fields (each in red), lumen phase field (blue).

### 2 Material and Methods

#### 2.1 Cell culture and sample preparation

##### 2.1.1 MDCK cell culture

Mardin-Darby Canine Kidney (MDCK) II cells were cultured using Gibco Minimum Essential Media (MEM) supplemented with 5% Fetal Bovine Serum (FBS). At 60-80% of confluency (every 2-3 days), cells are replated at a seeding density of $4.10^3$ cells.cm$^{-2}$. The MDCK cell lines used in this work with fluorescently labeled proteins are listed in Supp. Table 1.

| Cell line | Antibiotics | Origin |
|---|---|---|
| MDCK II WT | - | Yonemura Group [67] |
| MDCK II Actin GFP | 300 µg/ml Geneticin | Wedlich-Soldner group [68] |
| MDCK II MRLC GFP | 400 µg/ml Geneticin | Yonemura Group [67] |

Supp. Table 1: ***MDCK II cell lines used in this work.***

#### 2.2 Image analysis

The fluorescence intensity signals and the cell morphological characteristics were manually acquired using a customised Fiji [69] Plugin. Data were analysed using customised Python codes.

We give here details about the key protocols used to measure some features such as the measurement protocols of the cortex width, the cortex curvature, normalisation protocols for computing intensity ratios.

##### 2.2.1 Choice of the cell midplane for analysis

For each of the cell selected for analysis in the midplane of a cyst, we chose to perform the analysis in the middle plane of this cell as depicted in the scheme of Supp. Fig. 1.

##### 2.2.2 Estimate of the cortex width

To estimate the cortex width, we manually measured the intensity profile of the F-actin intensity signal along a line perpendicular to either the apical, basal or lateral cortices while averaging the line on a band of 10 pixels. Then, we normalised and we centered the intensity profile signal by :

$$I_{\text{norm}}^{\text{profile}} = \frac{I_{\text{profile}} - \langle I_{\text{background}} \rangle}{max(I_{\text{profile}})} \tag{1}$$

Then, we fitted the normalised and centralised profile by a gaussian intensity profile:

$$I_{\text{fit}}^{\text{profile}} = A \exp\left(\frac{-(x-\mu)^2}{2\sigma^2}\right) + B \tag{2}$$

with $A$, $B$, $\mu$, $\sigma$ constants determined by a non-linear least squares algorithm used to fit the previous the function.

The width of the cortex is then estimated to be the full width at half maximum of the gaussian fitted curve and is thus given by:

$$w = 2\sqrt{2\ln(2)}\sigma \tag{3}$$

To estimate the lateral cortex width, we divided by 2 the previous estimate since the F-actin signal at this location takes into account the width of the two neighbouring cells. These results are represented in Appendix Fig. 2(d) for the apical, lateral and basal cortices.

###### 2.2.3 Estimate of the cortex curvature

To estimate the cortex curvature we used the method of best fitting circle using a linear square method to fit a set of points [70] to obtain the radius of the circle best fitting a set of points that we manually drew on apical, basal or lateral cortices. The inverse of the radius of this circle was taken as the curvature of the cortex represented in Appendix Fig.2.

###### 2.2.4 Normalisation of the cortical intensity

We measured the total intensity and the number of pixels in segmented lines of 10 pixels thickness for apical, lateral and basal cortices with the help of a customised Fiji plugin. We normalised the signal intensity with the background intensity signal value as follows:

$$I_{\text{norm}} = \frac{I_{\text{ROI}} - N_{\text{pix}} I_{\text{background}}}{L} \tag{4}$$

where $I_{\text{ROI}} = \sum_{k=1}^{N_{\text{pix}}} I(i)$, $N_{\text{pix}}$ is the number of pixels in the ROI where the signal was measured and $L$ is the length of the part of the ROI. We used this normalisation in order to define a normalised intensity as an actual measure of the cortex density.

Then the intensity ratios are ratios of the normalised intensity at different locations of the cortex. Histograms of intensity ratios between different parts of the cortex for the F-actin and MRLC signals are represented in Appendix Fig. 2(b,c).

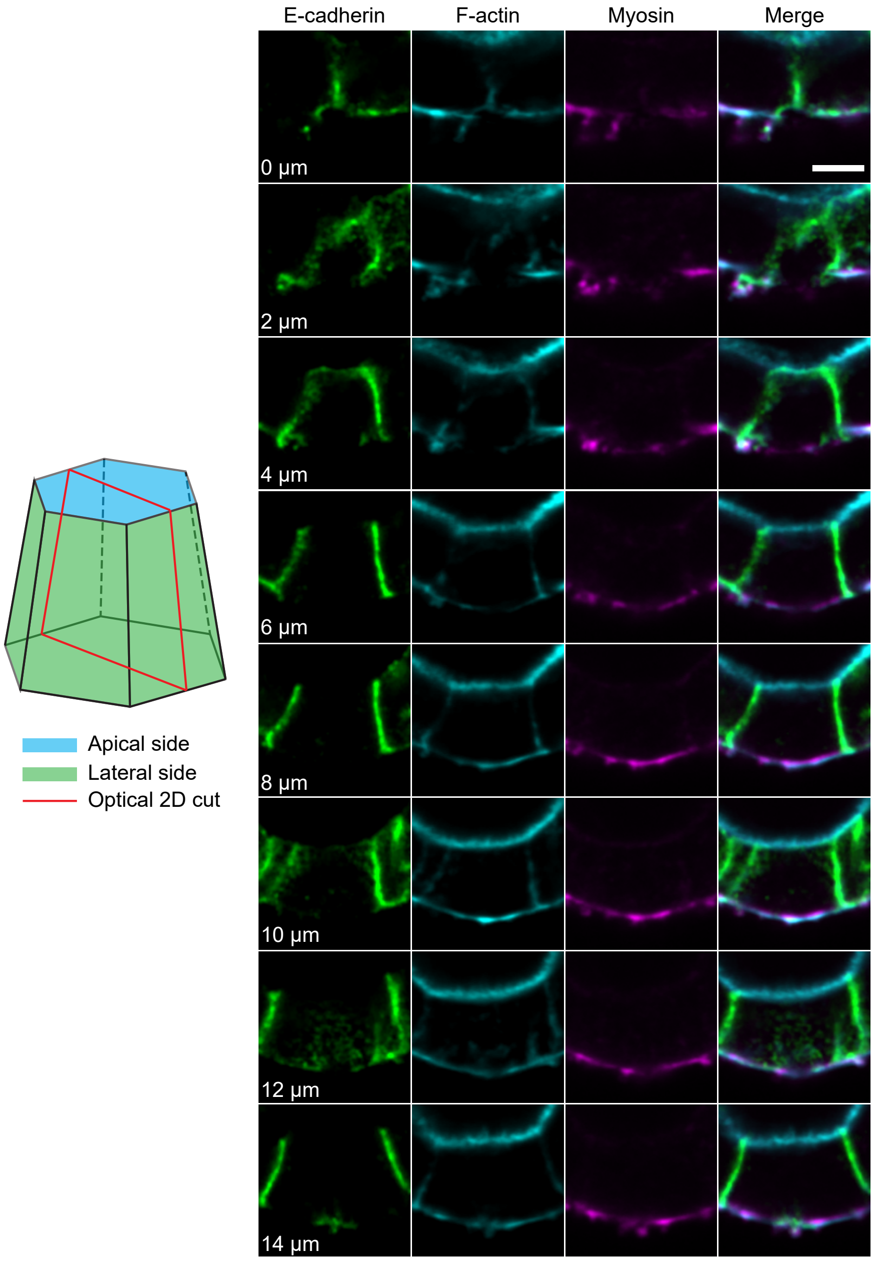

Supp. Fig. 1: ***2D cuts of 3D cell volumes reliably represent the acto-myosin distribution*** *- (Left) Scheme representing the optical slice (red frame) selected to perform quantifications. (Right) Snapshots of various 2D optical cuts of a cell in a MDCK cyst fixed and stained for F-actin (phalloidin, cyan), MRLC (magenta) and E-cadherin (green) at different positions along the microscope z-axis. Scale bar: 5µm.*

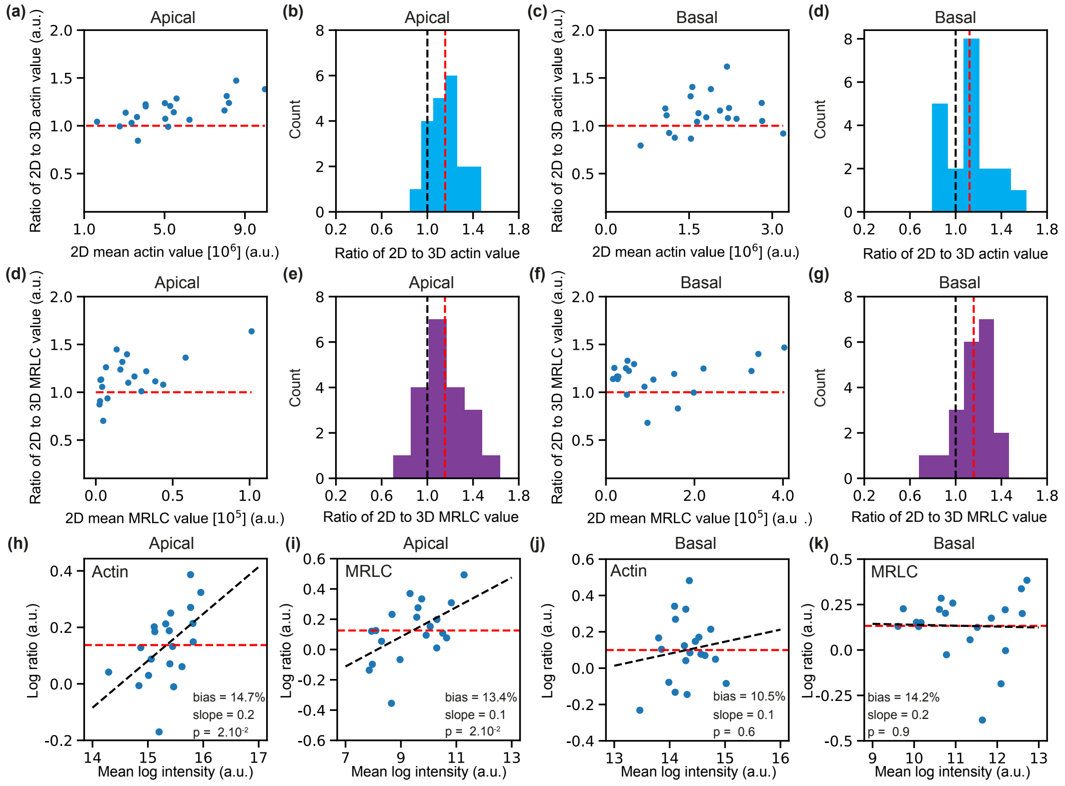

Supp. Fig. 2: ***Comparison between 3D mean values and 2D intersection.*** *- (a,c,e,g) Signal intensity value averaged along several sections as a function of signal intensity value in the midplane of the cell. (a) apical actin, (c) basal actin, (e) apical myosin, (g) basal myosin. (b,d,f,h) Histograms showing distributions across the z-axis. The horizontal axis is normalized by the 2D mean value, i.e., mean signal intensity value in the midplane. (a.u.) - red dashed line : mean of the distribution - black dashed line 0.0 value. (b) apical actin, (d) basal actin, (f) apical myosin, (g) basal myosin. (h-k) Bland-Altman tests to assess for fixed bias and proportional bias for F-actin and MRLC measurements at the apical and basal cortices. Bias, proportional bias (slope) and associated p-values are displayed on the graph themselves.*

### 3 Additional data and quantifications

We provide additional measurements that describe further the experimental system and other results mentioned in the main text.

#### 3.1 Cell shape diagram and correlations with other cell morphological quantities

##### 3.1.1 Lumen occupancy

We have quantified how distributed the cell shape parameter $\alpha$ is w.r.t its geometrical counterpart $\alpha_{\text{geometrical}} = R_\ell / R_s$, defined by assuming a unique cell shape within a cyst (Supp. Fig. 3). This graph shows that $\alpha$ is adapted to fairly capture the heterogeneity in cell shape.

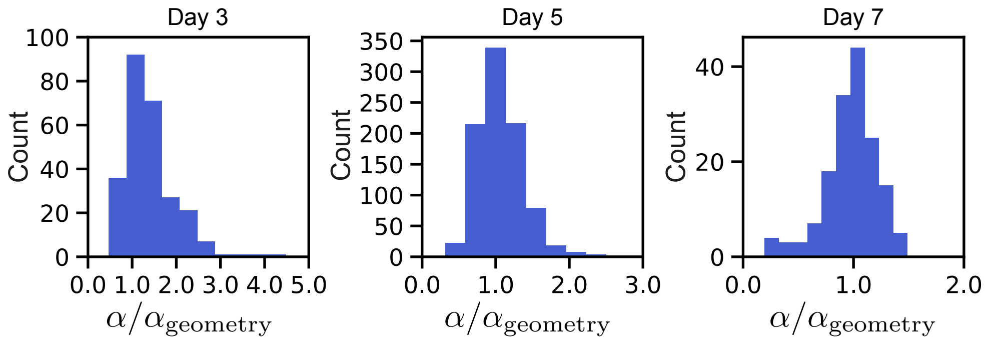

Supp. Fig. 3: ***Correlation between cell shape and lumen occupancy in MDCK cysts.*** - *Apical curvature correlation with cell shape. (d) Basal curvature correlation with cell shape. Distribution of the cell shape parameter* $\alpha = L_{ap}/L_{bas}$ *w.r.t* $\alpha_{geometry} = R_\ell/R_s$ *at day3, day 5 and day 7. For all measurements,* $N = 3$ *experiments, (middle)* $n = 791$ *cells; (left)* $n = 258$ *cells; (right)* $n = 158$ *cells.*

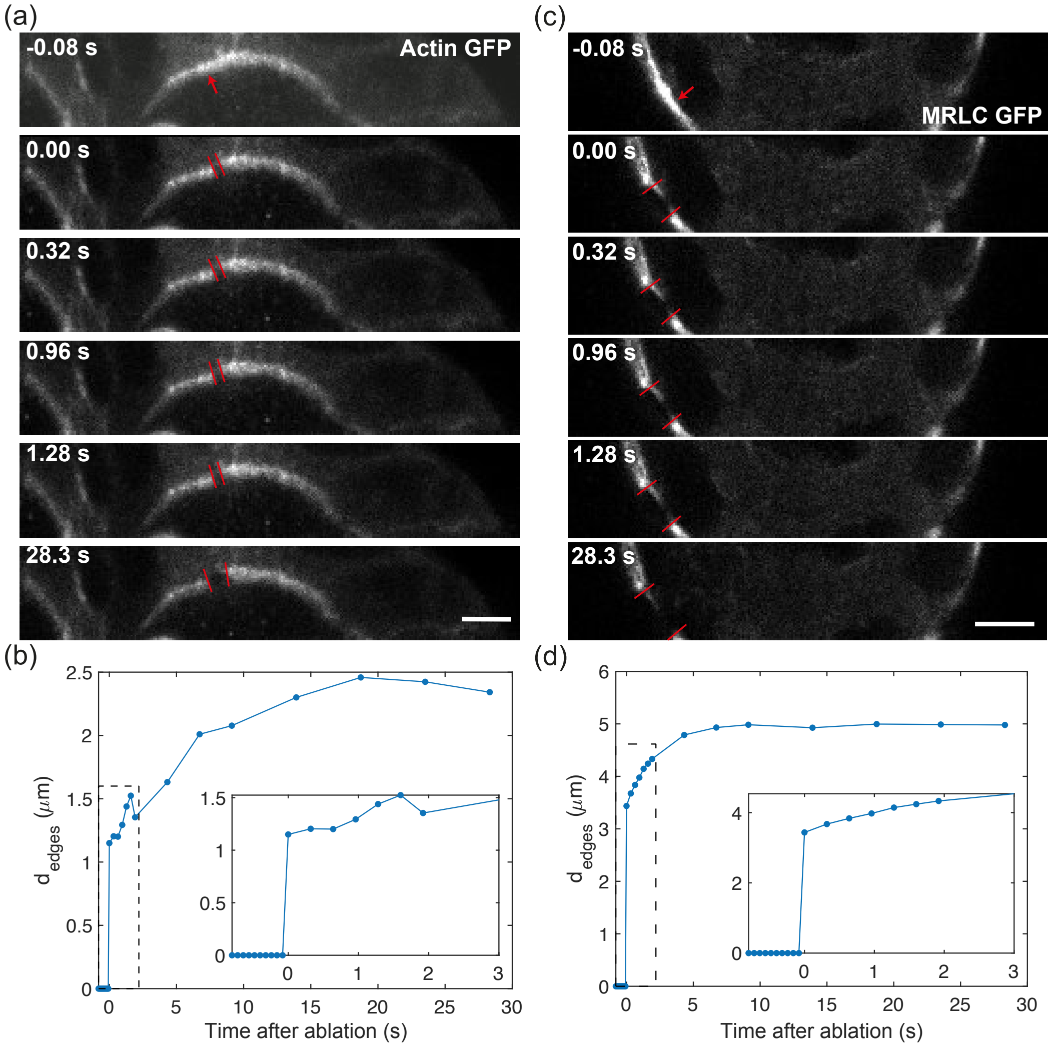

Supp. Fig. 4: ***Laser ablation experiments with improved time resolution.*** - *(a-c) Timecourse of laser ablation in the apical (a) and basal (c) cortex. Sites of ablation are highlighted with a red arrow, edges of the ablated cortex are highlighted with red bars; (b) Distance between the edges of the ablated cortex* $d_{edges}$ *as a function of time after laser ablation for apical (b) and basal (d) cortices. Scale bar is 5* $\mu m$*.*

#### 3.2 Laser ablation experiments

We detail here the additional results shown for the laser ablation experiments and their interpretation in term of inference of cortical stresses. We performed laser ablation experiments [26] to test the contractile or extensile nature of the active stresses generated by actin and myosin within the different parts of the cell cortex. When ablating apical, basal or lateral cortices, we observed an opening of the actin cortex of a few micrometers within seconds (Fig.5(c-d), Supp. Fig. 4). These openings were larger in the case of the basal cortex (Video 4, Fig. 2(e) and Fig. 5(c)) than in the case of the apical (Video 3, Fig. 2(e) and Fig. 5(b)) or the lateral (Video 5, Fig. 2(e) and Fig. 5(d)) cortices. This demonstrates that the cortices are more tensile at the basal sides rather than at the apical and lateral sides. We quantified the opening velocity $v_{\text{recoil}}^{\text{ini}}$ after laser ablation as a readout for comparing stress between the different apical, lateral and basal cortices, assuming they have similar damping coefficients [26].

#### 3.3 Perturbation experiments

We detail here the additional quantifications of the drug experiments targetting actin polymerization, myosin contractility and cell-cell adhesion. For the respective protocols, see section **??**.

##### 3.3.1 Depolymerizing F-actin

We incubated 5-day-old MDCK cysts with a depolymerizing actin drug LatA at a concentration of $c = 200$ nM and we monitored the change in cell shape up until 1 hour of incubation (Video 7). After 1h, we washed out LatA and changed back to fresh culture medium (see Material and Methods and Video 8). We quantified the changes in apical and basal curvatures (Appendix Fig. 7(b,c)) as well as the ratio between apical and basal length (Appendix Fig. 7(d)). Both apical and basal curvatures, as well as apical to basal length ratio increased in absolute values indicating that cells got deformed by a radial pushing force (*i.e* lumen hydrostatic pressure) following the depolymerisation of the actin cytoskleton upon LatA treatment.

In Supp. Fig.5, we show and quantify the change in cell shape following LatA washout (quantifications performed after 11 h of washout for a typical experiment). The washout of LatA induced a change in cell shape from an elongated cell to a more cuboidal one (Supp. Fig.5(b)). In this case, the actin ratio between the apical and basal cortices (Supp. Fig.5(c)), the ratio between the apical and basal cortical lengths (Supp. Fig.5(d)) and apical and basal curvatures (Supp. Fig.5(e-f)) came back to similar levels as prior LatA treatment.

Altogether, the washing out of LatA induced a polymerization of actin filaments in the actomyosin cortices and a transition back to the initial cellular shape within MDCK spheroids.

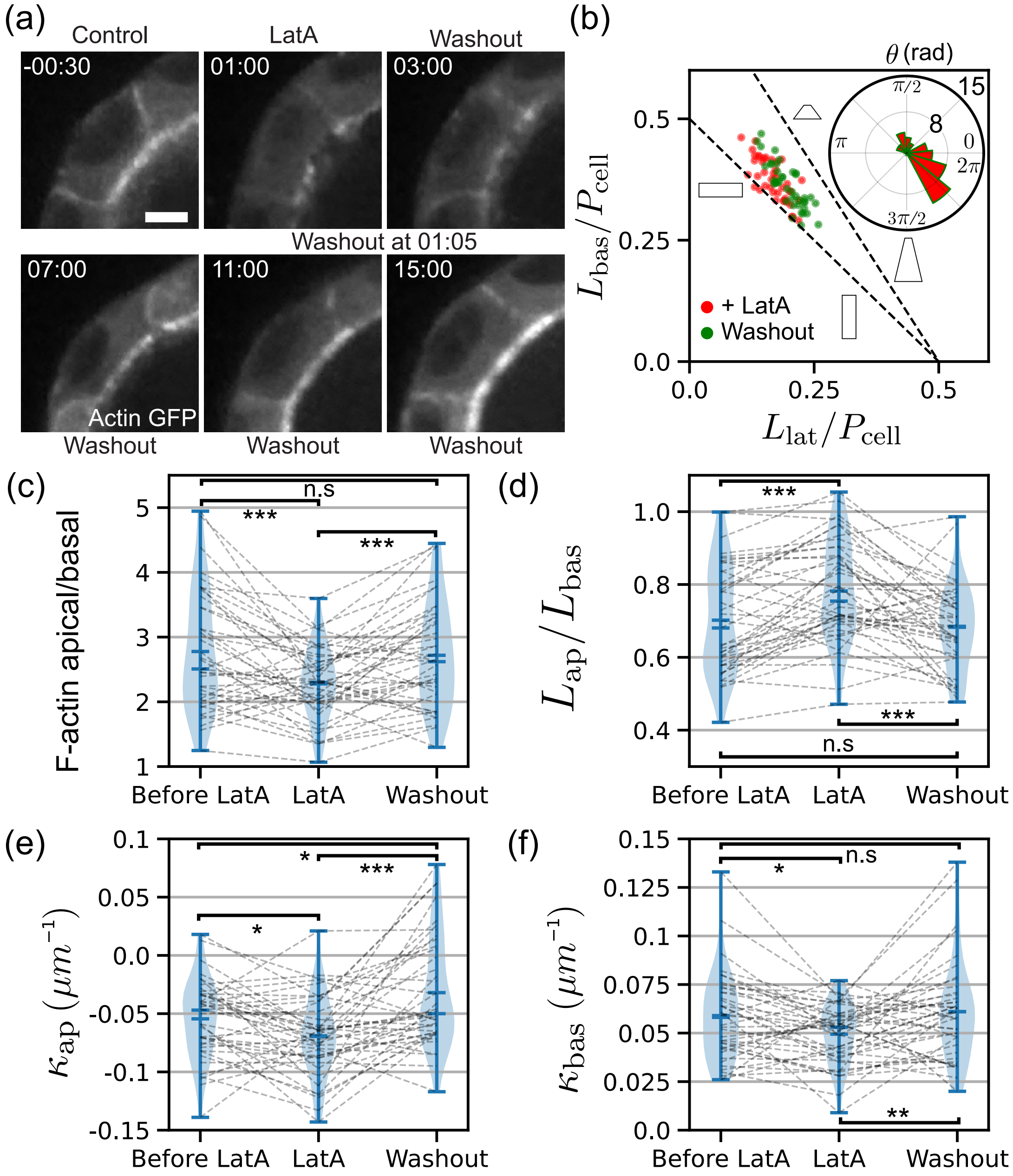

Supp. Fig. 5: ***Quantification of cell shape changes upon washout following a 200 nm LatA treatment.*** *(a) Inhibition of actin polymerisation by incubating MDCK cysts with LatA (c = 200 nM) following drug washout. Snapshots of an individual cell before treatment, after treatment and after washout. (Scale bar : 5 µm.) (b) Cell shape diagram showing the effect of LatA on cells: cells tend to become flatter and elongate longitudinally. Inset: distribution of θ, the angle of each trajectory vector in the cell shape diagram corresponding to Lat A washout. Quantification of changes in (c) apical to basal actin intensity ratios, (d) apical to basal length, (e) apical curvature and (f) basal curvature before and after LatA treatment and following washout. N = 3 biological repeats, typical quantification for N=1 and n = 41 cells.*

### 4 Multicellular phase field coupled to an active gel cortex model

Simulations were performed by adapting a phase field approach for multicellular systems [39,40,41] to the physics of active fluids [44,45]. Theoretical simulations of cysts were performed using a multicellular phase field model with lumen [40,41], additionally describing an extracellular elastic matrix around the cyst. We applied the resharpening method proposed in [42] and [43] which suppresses the inherent surface tension emerging from basic construction principle of the phase field model, to assure that the cellular surface tension comes solely from active gel and that the surface tension of lumen and ECM can be well controlled. The details of this theoretical model are given in the following subsections.

For our purpose, the phase-field approach offers a straightforward implementation of cell shape, lumen and extracellular matrix dynamics. This is why we chose the phase-field approach in this study. However, this does not limit the usage of the other approaches in future works. In general, each approach has inherent advantages compared with the other approaches, depending on the context; see Ref. [71] for more details.

#### 4.1 Multicellular phase-field

##### 4.1.1 Phase field variables to be implemented

In our multicellular phase-field model, organoids dynamics are simplified into three components: cells, lumen and ECM. Geometries of these components are represented by a corresponding phase field variable. We simulate $m$ cells by a phase field $\phi_i(\mathbf{r},t)$ ($i = 1, ..., m$, one for each cell), the lumen by the single phase field $\ell(\mathbf{r},t)$ and the ECM by $e(\mathbf{r},t)$. The regions of each phase field with 1 and 0 correspond to the inside and the outside of each component.

##### 4.1.2 Basic equations for dynamics

Although we also have implemented the "resharpening" (which will be explained in the next sub-sub-section) for the dynamics of phase fields, besides it, we essentially assumed that the dynamics is given by the gradient descent,

$$\tau_\phi \frac{\partial \phi_k}{\partial t} = -\frac{\delta E}{\delta \phi_k} \ , \quad \tau_\ell \frac{\partial \ell}{\partial t} = -\frac{\delta E}{\delta \ell} \ , \quad \tau_c \frac{\partial c}{\partial t} = -\frac{\delta E}{\partial c} \ , \tag{5}$$

with the free energy $E = E_\phi + E_\ell + E_c$, where each contribution (free energies of all cells $E_\phi$, the lumen $E_\ell$ and the ECM $E_c$) is given by

$$\begin{aligned} E_\phi = & \sum_k \int_\Omega \left[ \frac{D_\phi}{2} |\boldsymbol{\nabla} \phi_k|^2 + \frac{1}{4}\phi_k^2(1-\phi_k^2) \right] \mathrm{d}\mathbf{r} + \sum_k \frac{\alpha_\phi}{12} \left( V - \int_\Omega h(\phi_k) \mathrm{d}\mathbf{r} \right) \\ & + \sum_k \sum_{k' \neq k} \frac{\beta_\phi}{12} \int_\Omega h(\phi_k) h(\phi_{k'}) \mathrm{d}\mathbf{r} + \sum_k \sum_{k' \neq k} \frac{\eta_\phi}{12} \int_\Omega \boldsymbol{\nabla} h(\phi_k) \cdot \boldsymbol{\nabla} h(\phi_{k'}) \mathrm{d}\mathbf{r} \ , \end{aligned} \tag{6}$$

$$E_\ell = \int_\Omega \left[ \frac{D_\ell}{2} |\boldsymbol{\nabla} \ell|^2 + \frac{1}{4}\ell^2(1-\ell^2) \right] \mathrm{d}\mathbf{r} - \frac{\xi}{6} \int_\Omega h(\ell) \mathrm{d}\mathbf{r} + \sum_k \frac{\beta_{\ell\phi}}{12} \int_\Omega h(\ell) h(\phi_k) \mathrm{d}\mathbf{r} \tag{7}$$

and

$$
\begin{aligned}
E_c = & \int_\Omega \left[\frac{D_c}{2}|\boldsymbol{\nabla} c|^2 + \frac{1}{4}c^2(1-c^2)\right]\mathrm{d}\mathbf{r} - \frac{\xi_c}{6}\int_\Omega h(c)\mathrm{d}\mathbf{r} + \frac{\alpha_c}{12}\left(V_c - \int_\Omega h(c)\mathrm{d}\mathbf{r}\right) \\
& + \sum_k \frac{\beta_{c\phi}}{12}\int_\Omega h(c)h(\phi_k)\mathrm{d}\mathbf{r} + \frac{\beta_{c\ell}}{12}\int_\Omega h(c)h(\ell)\mathrm{d}\mathbf{r} + \sum_k \frac{\eta_{c\phi}}{6}\int_\Omega \boldsymbol{\nabla} h(c)\cdot\boldsymbol{\nabla} h(\phi_k)\mathrm{d}\mathbf{r}\ ,
\end{aligned}
\tag{8}
$$

respectively. Here, $D_\phi$, $D_\ell$, $D_c$ are diffusion constants also setting the width of the interface, $\alpha_\phi$ and $\alpha_c$ are bulk modulus of the cells and the ECM. $\beta_\phi$, $\beta_{\ell\phi}$, $\beta_{c\phi}$ and $\beta_{c\ell}$ are the coefficients controlling the exclusion volume interaction between cells, between cells and lumen, between cells and ECM and between lumen and ECM. $\eta_\phi$, $\eta_{c\phi}$ are coefficients for cell-cell adhesion and cell-ECM adhesion interactions. $\xi$ and $\xi_c$ are coefficent controlling lumen pressure and ECM pressure. $V$ and $V_c$ are target volumes of cell and ECM respectively. The function $h(x) = x^2(3-2x)$ is a monotonic function helpful for computation purposes. $\Omega$ is the spatial domain on which we aim to solve these equations.

Combining Eqs. (5)-(8) gives the time evolution equations for cells, lumen and ECM, as

$$
\tau_\phi \frac{\partial \phi_k}{\partial t} = D_\phi \boldsymbol{\nabla}^2\phi_k + \phi_k(1-\phi_k)\left(\phi_k - \frac{1}{2} + f_\phi\right)\ , \tag{9}
$$

$$
\tau_\ell \frac{\partial \ell}{\partial t} = D_\ell \boldsymbol{\nabla}^2\ell + \ell(1-\ell)\left(\ell - \frac{1}{2} + f_\ell\right)\ , \tag{10}
$$

and

$$
\tau_c \frac{\partial c}{\partial t} = D_c \boldsymbol{\nabla}^2 c + c(1-c)\left(c - \frac{1}{2} + f_c\right)\ , \tag{11}
$$

respectively, where

$$
\begin{aligned}
f_\phi(\psi,\phi_k,\ell,c) = & \ \alpha_\phi\left(V - \int_\Omega h(\phi_k)\mathrm{d}\mathbf{r}\right) & \text{[Cell volume control]} \\
& - \beta_\phi(\psi - h(\phi_k)) - \beta_{\ell\phi}h(\ell) - \beta_{c\phi}h(c) & \text{[Volume Exclusions]} \\
& + \eta_\phi\boldsymbol{\nabla}^2(\psi - h(\phi_k)) + \eta_{c\phi}\boldsymbol{\nabla}^2 h(c) & \text{[Adhesion terms]}
\end{aligned}
\tag{12}
$$

with $\psi = \sum_k \phi_k$,

$$
f_\ell(\psi, c) = -\beta_{\ell\phi}\psi - \beta_{c\ell}h(c) + \xi \tag{13}
$$

and

$$
f_c(\psi,\ell,c) = \alpha_c\left(V_c - \int_\Omega h(c)\mathrm{d}\mathbf{r}\right) - \beta_{c\phi}\psi - \beta_{c\ell}h(\ell) + \eta_{c\phi}\boldsymbol{\nabla}^2\psi + \xi_c\ . \tag{14}
$$

Here, $\tau_\phi$, $\tau_\ell$ and $\tau_c$ are characteristic timescales for phase field time evolution. In this work we considered $\tau_\phi = \tau_\ell = \tau_c = 1.0$.

###### 4.1.3 Resharpening of the phase field interface

In this work, instead of straightforwardly solving Eq. (9), (10) and (11), we applied the algorithm called 'resharpening' of the phase field interface, developed by Olsson [42] and Badillo [43].

In our case, this algorithm means the following modifications of the equations. When we compute each equation governing the dynamics of phase-field variables (Eqs.9, 10, 11), which had the general form

$$\tau \frac{\partial \phi}{\partial t} = D \boldsymbol{\nabla}^2 \phi + \phi(1-\phi)\left(\phi - \frac{1}{2} + f\right) , \tag{15}$$

we separate it into two steps:

$$\tau \frac{\partial \phi}{\partial t} = \phi(1-\phi) f \tag{16}$$

$$\tau \frac{\partial \phi}{\partial t^*} = \boldsymbol{\nabla} \cdot \left( D \boldsymbol{\nabla} \phi - \sqrt{2D}\phi(1-\phi)\frac{\boldsymbol{\nabla}\phi}{|\boldsymbol{\nabla}\phi|}\right) . \tag{17}$$

Equation (16) computes the phase field changes according to interactions with other phase fields. Equation (17) is used to control the phase field interface profile. Operationally, after the phase $\phi$ is integrated using Eq. (16) for a timestep $\Delta t$, $\phi$ is computed in an auxillary time $t^*$ according to the conservative equation (17) until the phase field converges ($\partial\phi/\partial t^* = 0$). See Refs. [42,43] for more details.

The effects of this modification of equations can be listed as follows: (i) Along the direction perpendicular to the interface, each phase-field variable keeps the steady-state profile of Eq. (17), which is the hyperbolic tangent $\phi(r) = [1 + \tanh(r/w)]/2$ with the fixed interface width $w = \sqrt{D}$. (ii) Since Eq. (16) itself does not include the diffusion term, the built-in surface tension due to the diffusion terms is eliminated. Otherwise, reflecting the fact that diffusion terms come from the terms with $D$'s in the free energies [Eqs. (6)-(8)], they inevitably introduce surface tension for finite values of $D$. Note that the effect (i) is beneficial when we confine the active gel into the phase-field interface later. The effect (ii) lets the active gel solely provide surface tension to the cell as well as allows us to better control the surface tension of lumen and ECM.

#### 4.2 Coupling Active gel model of the cortex with the phase fields

##### 4.2.1 Active gel equations for the cortex

We applied the resharpening method proposed in [42] and [43] which suppresses the inherent surface tension emerging from basic construction principle of the phase field model, to assure that the cellular surface tension comes solely from active gel and that the surface tension of lumen and ECM can be well controlled.

To model the cortex using the active gel theory, we use an extension of the 1D model developed in [44] that we adapt from the work of Levernier and Kruse [45].The active gel is described with a continuum model of its density $\rho$. It evolves accordingly to a conservation equation ($\alpha$, $\beta$ are the $x$, $y$ coordinates):

$$\partial_t \rho + \boldsymbol{\nabla} \cdot (\rho \boldsymbol{v}) = -A_{\text{deg}}\rho + A_{\text{gen}}|\boldsymbol{\nabla}\phi|\delta\left(\phi - \frac{1}{2}\right) \tag{18}$$

The L.H.S of Eq.18 is the time derivative of the gel density and the divergence of the flux of gel. The R.H.S of this equation relates to the turnover of the active gel: the first term is the degeneration of the gel assumed to be isotropic, the second term is the source term, which we assume to be restricted in the cell cortex. Here and after, in practice, we approximate the Dirac's delta by the normal distribution with the finite width $\epsilon$;

$$\delta\left(\phi(\mathbf{r})-\frac{1}{2}\right)\sim\frac{1}{\sqrt{2\pi}\epsilon}\exp\left(-\frac{\phi-1/2}{2\epsilon^2}\right) . \tag{19}$$

The velocity of the gel can be derived after solving the force balance equation for the gel:

$$\partial_\beta\sigma_{\alpha\beta}=\gamma v_\alpha , \tag{20}$$

where $\sigma_{\alpha\beta}$ is the total stress tensor, $v_{\alpha\beta}=(\partial_\alpha v_\beta+\partial_\beta v_\alpha)/2$ is the velocity gradient tensor and $\gamma$ a friction coefficient. The stress tensor writes as:

$$\sigma_{\alpha\beta}=\sigma_{\text{viscous}}+\sigma_{\text{active}}=2\eta v_{\alpha\beta}-\Pi_{\alpha\beta}(\rho) \tag{21}$$

This tensor $\Pi_{\alpha\beta}$ is a part of the stress tensor independent of the velocity field, and we assume that it has two contributions, an osmotic effective pressure part and an active stress part. We choose to follow [44] and [45] to express $\Pi$ in terms of the gel density $\rho$. To guarantee the symmetry, we give the definition of $\Pi(\rho)$ in the reference frame of tangential and orthogonal directions of the cell cortex:

$$\Pi(\rho)=\begin{bmatrix}\Pi_\perp & 0\\ 0 & \Pi_{\|}\end{bmatrix} , \tag{22}$$

where

$$\Pi_\perp=-a_\perp\rho^3+b\rho^4 , \ \Pi_{\|}=-a_{\|}\rho^3+b\rho^4 . \tag{23}$$

Here, for $a_{\perp/\|}>0$ we reflect the active contractile stresses, if $a_{\perp/\|}<0$ the active stresses are extensile. One can view the contributions to the third power of the gel density as a translation of the microscopic interaction between actin filaments and myosin motors in which a single motor ($\rho$) needs to interact with two actin filaments ($\rho^2$) in order to generate a stress. The contractility of motors is capable to induce a phase separation in the gel [44].

To transfer back Eq. (22) in the cartesian coordinate system, we use the definition of the normal vector to the phase field interface, $\hat{\mathbf{n}}=(\boldsymbol{\nabla}\phi)/|\boldsymbol{\nabla}\phi|$, and apply a rotation matrix to $\Pi$ given in Eq. (22), which gives an expression of the active tensor as:

$$\Pi(\rho)=\begin{bmatrix}\Pi_{xx} & \Pi_{xy}\\ \Pi_{yx} & \Pi_{yy}\end{bmatrix}=\begin{bmatrix}-a_{\|}\rho^3+b\rho^4+(-a_\perp+a_{\|})\rho^3n_x^2 & (-a_\perp+a_{\|})\rho^3n_xn_y\\ (-a_\perp+a_{\|})\rho^3n_xn_y & -a_{\|}\rho^3+b\rho^4+(-a_\perp+a_{\|})\rho^3n_y^2\end{bmatrix} \tag{24}$$

Solving the dynamical equations for the active gel density can describe many different behaviours. For high enough tangential contractile active stresses, spontaneous protrusive active gel accumulation appears. It is also possible to find a parameter regime in which the gel has a stable step profile going away from the cell cortex [45].

It is to be noted that, for Eq. (23), the choice of the functional form of $\Pi$ is not essential. Another choice can be made, for instance $\Pi_{\perp/\|}(\rho)=-a_{\perp/\|}\rho+b\rho^2$, as long as it reflects that the passive osmotic pressure always dominates at large densities and in the case where activity of motors is large enough, the effective osmotic pressure can become a non-monotonic function of density.

###### 4.2.2 Confinement of the active cortex into the cell phase-field interface

The coupling term between the active gel and the phase-field of the cell is encoded in an additional free energy term for each cell:

$$F_{\text{coupling}}[\phi, \rho] = \underbrace{-\int_{\Omega} f_{\text{coup}} \rho |\boldsymbol{\nabla} \phi(\boldsymbol{r})| \delta \left( \phi(\boldsymbol{r}) - \frac{1}{2} \right) \mathrm{d}\boldsymbol{r}}_{\text{confinement of the gel to the cortex}} \tag{25}$$

where $f_{\text{coup}}$ coefficient controls the strength of coupling, we set it to $f_{\text{coup}} = 1.0$ in what follows. This term in the free energy provides a new term in the evolution equation of the cell phase-field $\phi$ [Eq. (9)]:

$$\tau_{\phi} \frac{\partial \phi}{\partial t} = -\frac{\delta E_{\phi}}{\delta \phi} - \frac{\delta F_{\text{coupling}}[\phi, \rho]}{\delta \phi} \tag{26}$$

and we can compute that:

$$-\frac{\delta F_{\text{coupling}}[\phi, \rho]}{\delta \phi} = -f_{\text{coup}} \delta \left( \phi - \frac{1}{2} \right) \boldsymbol{\nabla} \cdot (\rho \hat{\boldsymbol{n}}) \tag{27}$$

with $\hat{\boldsymbol{n}} = \frac{\boldsymbol{\nabla} \phi}{|\nabla \phi|}$, the normal vector to the phase-field $\phi$. Then we see that the effect of this coupling term is to move the phase-field interface boundary in the direction opposite to the normal vector. The magnitude of the displacement is relative to the density of the gel $\rho$. For an uniform cortex of constant density, the coupling term induces a global contractility which counteracts the volume control and adhesions already set in the cell shape dynamics.

In practice, one can write $\boldsymbol{\nabla} \cdot (\rho \hat{\boldsymbol{n}}) = \hat{\boldsymbol{n}} \cdot \boldsymbol{\nabla} \rho + \rho \boldsymbol{\nabla} \cdot \hat{\boldsymbol{n}}$, and for numerical stability reasons, we neglected the second term, $\rho \boldsymbol{\nabla} \cdot \hat{\boldsymbol{n}}$. This approximation can be justified when the interface width $w = \sqrt{D}$ is much smaller than the inverse of the local curvature of the interface $C$ because $|\hat{\boldsymbol{n}} \cdot \nabla \rho| \sim O(\rho^1 w^{-1} C^0)$ whereas $|\rho \boldsymbol{\nabla} \cdot \hat{\boldsymbol{n}}| \sim O(\rho^1 w^0 C^1)$. Since we will use only the small $D$ in this work, this is indeed the case.

##### 4.3 Numerical implementation

###### 4.3.1 Summary of the equations and calculation steps

We integrated the equations in the following order:

- the active gel dynamics was solved for a timestep $\Delta t$ with the values of the cell phase-fields considered at time $t$. To this end, we integrated Eq. 18 using a finite difference method in the Fourier space, see sub-sub-section 4.3.2 for details of the procedure.
- Eq.16 for all the phase-field variables $\phi_k$ (for each cell $k$), $\ell$ and $c$ were updated for a timestep $\Delta t$ with the values of the active gel density considered at time $t$.
- Resharpening step given by Eq.17 for each cell $\phi_k$, lumen $\ell$ and ECM $c$ phase-field interfaces.

For simulations of multicellular phase-field, we used a custom written C code including OpenACC directives implemented with the PGI compiler version 20.4 [72] to perform GPU accelerated computations.

Although we are focusing in the present study on 2D implementation of the previous set of equations, it is in principle possible to extend its implementation to 3D. Indeed, the multicellular phase-field model has already been successfully implemented in three dimensions [39,40,41]. However, in practice, the implementation of the active gel model and its coupling with the phase-field model may require a careful numerical scheme for 3D implementation. Therefore, we have decided not to include the actual 3D calculations in this study.

##### 4.3.2 Details of the numerical schemes

We discretised the set of equations on the simulation grid and we used periodic boundary conditions for all the phase fields and active gel density. We used finite difference methods on the regular-square lattice to compute the quantities at $t + \Delta t$. For phase fields and active gels, we used a forward difference for time integration and we defined laplacians with a 8 point stencil method.

To compute the force balance of the active gel, we computed $\rho\boldsymbol{v}$ using Fourier transform. Shortly, the process is as follows. The force balance equation is written as

$$\partial_\beta \sigma_{\alpha\beta} = \gamma v_\alpha \ . \tag{28}$$

Since $\sigma_{xx} = \eta\partial_x v_x - \Pi_{xx}$, $\sigma_{xx} = \eta\partial_y v_y - \Pi_{yy}$ and $\sigma_{xy} = \sigma_{yx} = \frac{1}{2}\eta(\partial_x v_y + \partial_y v_x) - \Pi_{xy}$, we have

$$\begin{aligned} 2\eta\partial_{xx}v_x - \partial_x\Pi_{xx} + \eta(\partial_{yy}v_x + \partial_{yx}v_y) - \partial_y\Pi_{xy} = \gamma v_x \ , \\ \eta(\partial_{xx}v_y + \partial_{xy}v_x) - \partial_x\Pi_{xy} + 2\eta\partial_{yy}v_y - \partial_y\Pi_{yy} = \gamma v_y \ . \end{aligned} \tag{29}$$

We then take the Fourier transform of each equation. We label with a $\tilde{}$ all Fourier transformed functions such as $FT[v_x] = \tilde{v}_x$. In Fourier space, the differential operator $\partial_x$ corresponds to multiplying by $ik_x$ (where $k_x$ are spatial frequencies equally spaced of $2\pi/N$, where $N$ is the extent of the grid) and we have:

$$\begin{aligned} -2\eta k_x^2\tilde{v}_x - ik_x\tilde{\Pi}_{xx} + \eta(-k_y^2\tilde{v}_x - k_xk_y\tilde{v}_y) - ik_y\tilde{\Pi}_{xy} = \gamma\tilde{v}_x \\ \eta(-k_y^2\tilde{v}_y - k_xk_y\tilde{v}_x) - ik_x\tilde{\Pi}_{xy} - 2\eta k_y^2\tilde{v}_y - ik_y\tilde{\Pi}_{yy} = \gamma\tilde{v}_y \end{aligned} \tag{30}$$

These Fourier transforms of Eqs.29 allow to compute $\tilde{v}_x$ and $\tilde{v}_y$. We can then take these quantity back to real space to compute $\rho\boldsymbol{v}$. Then we compute $\tilde{\rho\boldsymbol{v}}$, and we update $\tilde{\rho}$ at $t + \Delta t$ with Eq. (18) using a finite difference method in the Fourier space before transforming back $\tilde{\rho}$ to the real space.

##### 4.3.3 Initial conditions

We started each simulation by putting $N_{\text{cells}}$ circular cells of radius $R$ equally placed on a best fitting circle of radius $R_{\text{cyst}}^{\text{ini}} = 0.9N_{\text{cells}}R/\pi$ so cells overlap initially. $R_{\text{cyst}}$ is centered on $(X_c,\, Y_c)$.

The initial profile for cell phase field is defined by:

$$\phi(i,j) = \frac{1}{1 + \exp\left(\frac{\Delta x\sqrt{(i-x_c)^2+(j-y_c)^2} - R}{w}\right)} \tag{31}$$

where $(i,\, j)$ represents the grid coordinates, $(x_c,\, y_c)$ are the coordinates of the cell center, $R = 1.0$ its initial radius and $w$ as previously defined.

We defined the initial lumen and ECM phase fields by using the cell phase fields. All points in the area of $\Omega$ confined by the cell monolayer (*i.e.*, $\sqrt{(i - X_c)^2 + (j - Y_c)^2} < R_{\text{cyst}}$) are defined as being part of the lumen phase: there $\ell(i,j) = 1 - \sum_{k=1}^{N_{\text{cells}}} \phi_k(i,j)$ and $\ell = 0$ elsewhere. All the rest of $\Omega$ (*i.e.*, $\sqrt{(i-X_c)^2+(j-Y_c)^2} > R_{\mathrm{cyst}}$) is attributed to the ECM phase field with a value of $c(i,j) = 1 - \sum_{k=1}^{N_{\mathrm{cells}}} \phi_k(i,j)$ and $c(i,j) = 0$ elsewhere.

The initial active gel density is computed from the initial cell phase-field:

$$\rho(i,j) = \rho_0 \exp\left(\frac{1}{1+\sigma} - \frac{1}{4\phi^2(i,j)(3-2\phi(i,j))(1-\phi^2(i,j)(3-2\phi(i,j))+\sigma)}\right) \tag{32}$$

where $\rho_0$ is the initial active gel density, taken to be close to the equilibrium value set by the turnover dynamics and $\sigma = 0.01$ is a tolerance threshold for defining $\rho$. If $\rho(i,j) < \rho_0 \exp(-1.0/(2.0\sigma) + 1.0/(1.0+\sigma))$, we set $\rho(i,j) = 0.0$.

###### 4.3.4 Definition of the apical, basal and lateral regions

In this work, as described in the next subsubsection, we used different values of active-gel parameters ($A_{\mathrm{gen}}$, $A_{\mathrm{deg}}$, $a_\perp$, $a_\parallel$ and $b$) between the apical, basal and lateral regions. To define each label of the region, we took advantage of the phase-field formulation. For a cell phase-field $\phi_i$ and for each point in the corresponding domain $\Omega_{\mathrm{cells}}$, we find the maximum value among $[\phi_i\phi_j$ (for j in neighbours)$, \phi_i\ell, \phi_i c]$. Then, the entire domain is splitted into 'apical' domain (when $\phi_i\ell$ is the max value), 'basal' domain (when $\phi_i c$ is the max value) or 'lateral' domain (when any overlapping with the neighbouring cells makes $\phi_i\phi_j$ the max value).

###### 4.3.5 Parameters and other detailed settings used for each result

For the main simulations with $N = 6$ initial cells, we integrated the equations on grids of area $\Omega = 1200 \times 1200$, with a spatial resolution $\Delta x = \Delta y = 0.01$ and a time resolution of $\Delta t = 0.002$. In order to accelerate the computation, we integrated the phase field dynamics of each cell on a smaller domain of area $\Omega_{\mathrm{cells}} = 200 \times 200$, centered on the cell center of mass that is computed at every timestep.

For the phase-field dynamics, we always used $w = \sqrt{D} = 0.07$, $\alpha_\phi = 1.0$, $\alpha_c = 0.002$, $\beta_\phi = \beta_{\ell\phi} = \beta_{c\phi} = \beta_{c\ell} = 1.0$, $\tau_\phi = \tau_c = \tau_\ell = 1.0$, $\eta_\phi = 0.03$, $\eta_{c\phi} = 0.001$. We set the ECM pressure $\xi_c = 0.0$, $V_c = \Omega$ and $V = 3.15$. To couple the active gel to the cell interface, we used a $\delta$ function defined with $\epsilon = 0.2$ and a coupling coefficient $f_c = 1.0$.

For the active gel dynamics, we used various values of the parameters and various detailed settings. In this paragraph, we explain how we chose them for each result. The friction coefficient was fixed to $\gamma = 1.0$, viscosity coefficient was fixed to $\eta = 1.0$ and, depending on the specific simulations, the values of $a_\perp$, $a_\parallel$, $A_{\mathrm{gen}}$ and $A_{\mathrm{deg}}$ were changed for the different figures as follows:

- For Fig. 3(a), the initial condition was the steady-state obtained using the protocol described in section 4.3.3 for 6 cells, $\xi = 0.15$, $\eta_\phi = 0.03$, $\alpha_c = 0.002$ and the rest of the parameters listed above with $A_{\mathrm{gen}} = 1.0$ and $A_{\mathrm{deg}} = 100$ for apical, lateral and basal cortices. Video 6 is a movie showing how steady-state is reached. Next, for all simulations, we set symmetric cortices with $A_{\mathrm{deg}}^{\mathrm{apical}} = A_{\mathrm{deg}}^{\mathrm{basal}} = A_{\mathrm{deg}}^{\mathrm{lateral}} = 100.0$ and we indicated values of $A_{\mathrm{gen}}^{\mathrm{apical}} = A_{\mathrm{gen}}^{\mathrm{basal}} = A_{\mathrm{gen}}^{\mathrm{lateral}}$ on the panel of the figure. No additional active stresses were added for these simulations, in practice we removed the term $\partial_\alpha(\rho v_\alpha)$ from Eq. 18. Simulations were run for 300000 steps and the top left cell shape was analysed to display the results in Fig. 3(b).
- For Fig. 6(c), the initial condition was the same as in Fig. 3(a). Then asymmetric turnover rates ($A_{\mathrm{gen}}^{\mathrm{apical}}$, $A_{\mathrm{gen}}^{\mathrm{basal}}$, $A_{\mathrm{gen}}^{\mathrm{lateral}}$) were chosen as diplayed in the panel of Supp. Fig. 6(c). No

additional active stresses were added for these simulations, in practice we removed the term $\partial_\alpha(\rho v_\alpha)$ from Eq. 18. Simulations were run for 100000 steps and the top left cell shape was analysed to display the results in Supp. Fig. 6(d).

- For Fig. 3(c), the inital condition was the result of computing 700000 steps of the set of parameters described in Fig. 3(a) for $A_{\text{gen}}^{\text{apical}} = A_{\text{gen}}^{\text{basal}} = A_{\text{gen}}^{\text{lateral}} = 0.25$. Next, we added additional active stresses of different nature (extensile or contractile) by choosing coefficients $a_\perp$, $a_\parallel$ for the apical, basal and lateral cortices.

  The baseline parameters were chosen to be: $a_\perp^{\text{lateral}} = 500$, $a_\perp^{\text{apical}} = a_\perp^{\text{basal}} = 50000$, $a_\parallel^{\text{lateral}} = 1000$ and $b^{\text{apical}} = b^{\text{basal}} = b^{\text{lateral}} = 1.0$.

  Then we chose to explore four different configurations by alternating the signs of $a_\perp^{\text{apical}} = \pm 10^6$ and $a_\perp^{\text{basal}} = \pm 10^6$ as displayed in the panel of Fig. 3(c). Simulations were run for 300000 steps and the top left cell shape was analysed to display the results in Fig. 3(d).

- For Fig. 3(e), we chose parameters fitting the experimental characterisation that we performed (Fig. 2 and Table 1).

  From the FRAP experiments, we measured (Fig. 2 and Table 1):

$$\tau_{\text{apical}} = 4\tau_{\text{basal}} = 4\tau_{\text{lateral}} \tag{33}$$

  Because we probed the dynamics of assembly and dissassembly of the actin network with FRAP experiments, we translated these measurements into values of $A_{\text{deg}}$:

$$A_{\text{deg}}^{\text{apical}} = \frac{1}{4} A_{\text{deg}}^{\text{basal}} = \frac{1}{4} A_{\text{deg}}^{\text{basal}} \tag{34}$$

  From the actin intensity measurements, we measured (Fig. 1-2, Appendix Fig. 2(b,c)), assuming that intensities of fluorescently labelled actin is a readout for actin density:

$$\rho_{\text{actin}}^{\text{apical}} = 5\rho_{\text{actin}}^{\text{basal}} = 7\rho_{\text{actin}}^{\text{lateral}} \tag{35}$$

  To use this information, we made the hypothesis that the measurements reflected the steady-state density of the actin cortex. We could then write Eq. (18) at steady state (with no actin flows) and find that:

$$\frac{A_{\text{gen}}^{\text{apical}}}{A_{\text{deg}}^{\text{apical}}} = 5\frac{A_{\text{gen}}^{\text{basal}}}{A_{\text{deg}}^{\text{basal}}} = 7\frac{A_{\text{gen}}^{\text{lateral}}}{A_{\text{deg}}^{\text{lateral}}} \tag{36}$$

  By combining these two results we got:

$$A_{\text{gen}}^{\text{apical}} = \frac{5}{4} A_{\text{gen}}^{\text{basal}} = \frac{7}{4} A_{\text{gen}}^{\text{lateral}} \tag{37}$$

The laser ablation experiments provided insight into the cortical tension at apical, lateral and basal sides of the cells (Fig. 2 and Table 1). These measurements can serve as estimate for the active contractile stresses. The experiments provided ratios of contractile tensions through measurements of initial recoil velocity [26] :

$$\langle v_{\text{recoil}}^{ini,basal} \rangle = \frac{3}{2} \langle v_{\text{recoil}}^{ini,apical} \rangle \tag{38}$$

To estimate ratios of contractile stresses we made the hypothesis that the contractile stresses measured by laser ablation represented the tangential active stresses in the active gel model. Then we had from the definition of $\Pi(\rho)$:

$$a_{\parallel}^{\text{basal}} \left(\rho_{\text{actin}}^{\text{basal}}\right)^3 = \frac{3}{2} a_{\parallel}^{\text{apical}} \left(\rho_{\text{actin}}^{\text{apical}}\right)^3 \tag{39}$$

Using the previous measured actin densities, we estimated that:

$$a_{\parallel}^{\text{basal}} = \frac{375}{2} a_{\parallel}^{\text{apical}} \tag{40}$$

In practice, to generate the result shown in Fig. 3(e), we started with an initial condition of the steady state like in Fig. 3(a) and Supp. Fig. 6(c). Then, we looked for a steady state that satisfied the previous relationships between $A_{\text{gen}}^{\text{apical}}$, $A_{\text{gen}}^{\text{lateral}}$, $A_{\text{gen}}^{\text{basal}}$, $A_{\text{deg}}^{\text{apical}}$, $A_{\text{deg}}^{\text{lateral}}$, $A_{\text{deg}}^{\text{basal}}$ with $A_{\text{gen}}^{\text{apical}} = 1.0$ and $A_{\text{deg}}^{\text{apical}} = 100.0$. With the previous relationship between $a_{\perp}^{\text{apical}}$ and $a_{\perp}^{\text{basal}}$, we set $a_{\perp}^{\text{basal}} = 50000$, $a_{\perp}^{\text{lateral}} = 100$, $a_{\parallel}^{\text{lateral}} = 100$ and $b^{\text{apical}} = b^{\text{basal}} = b^{\text{lateral}} = 1.0$.

To reach steady-state, we had to explore the parameter space of lumen pressure and cell adhesion and we find the situation depicted in Fig. 3(e) by adjusting $\xi = 0.17$ and $\eta = 0.02$.

- For Fig. 4(e), we started from the steady state of Fig. 3(e), $\xi = 0.17$, $\eta = 0.02$ and the parameters deduced from the experiments as described above. Then we modified globally $A_{\text{gen}}$ as indicated on the panel of Fig. 4(g). Simulations were run for 250000 steps and the top left and top cell shapes were analysed to generate the plot in Fig. 4(f).

#### 4.4 Sensitivity analysis

To perform sensitivity analysis, we tried the one-at-a-time method in which we vary the value of a single parameter while keeping the other being constant, by selecting parameters $x$, and see how the values of a few selected variables $y$ at a given time step are changed. The initial condition for this analysis is the steady state shown in Fig. 3e, from which we run a simulation for 100000 numerical steps. Then, we measure the variables, and as a metric to quantify the sensitivity, compute the slope of the variation of the selected variable $y$ with respect to the change of the selected parameter $x$, in the parameter range $[x_{\min}, x_{\max}]$, defined as:

$$\Omega_y = \frac{y(x_{\max}) - y(x_{\min})}{x_{\max} - x_{\min}} .$$

Here, as the variables, we particularly focus on the cell shape parameter $y = \alpha$ and of lumen occupancy $y = L_{\text{occ}}$, and the results are summarized in Table 2. We also report our measurements in Supp. Fig. 6 for cell shape parameter $\alpha$ and in Supp. Fig. 7 for lumen occupancy $L_{\text{occ}}$ We found low sensitivity for $\epsilon$, $\eta_{c\phi}$ (adhesion between ECM and cells), $w$ (width of the cortex) and significant sensitivity for $\alpha_c$ (ECM elasticity), $\eta$ (cell-cell adhesion) and $\xi$ (lumen pressure).

Moreover, it is worth noting that $\beta$, an artificial parameter to implement the volume exclusion in our framework, appropriately shows low sensitivity. This insensitivity is, in fact, a desirable feature and justifies our choice to fix the $\beta$ value at a single value in our study. Regarding parameters that control the active gel dynamics, we see low sensitivity around the steady-state of Fig. 3e for $a^{\text{apical/basal/lateral}}$ and $a_{\parallel}^{\text{apical/basal/lateral}}$ which we also discuss in Appendix Fig. 8 and in Appendix 4(b).

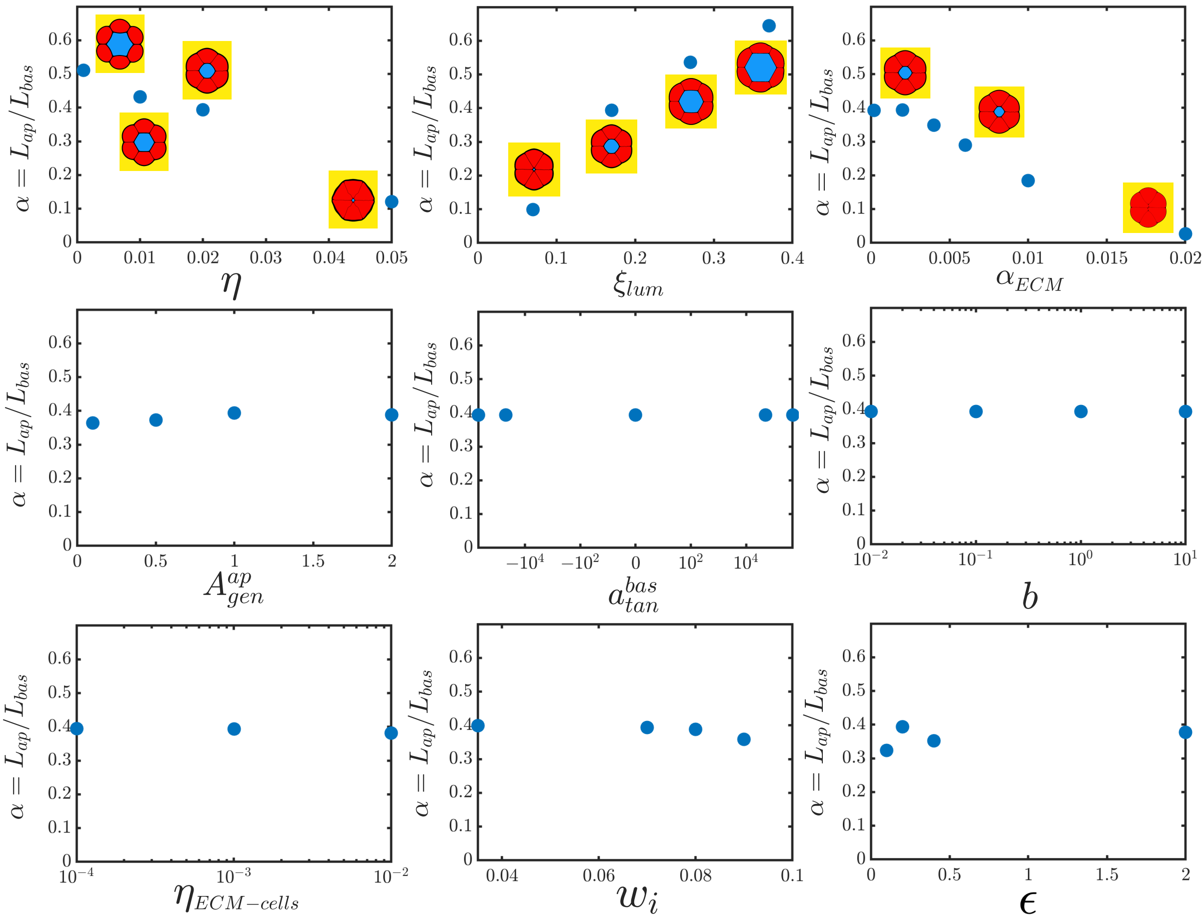

Supp. Fig. 6: ***Sensitivity analysis for the cell shape parameter*** $\alpha$***.*** *Sensitivity analysis. Cell shape parameter* $\alpha$ *is measured as one parameter value is modified.*

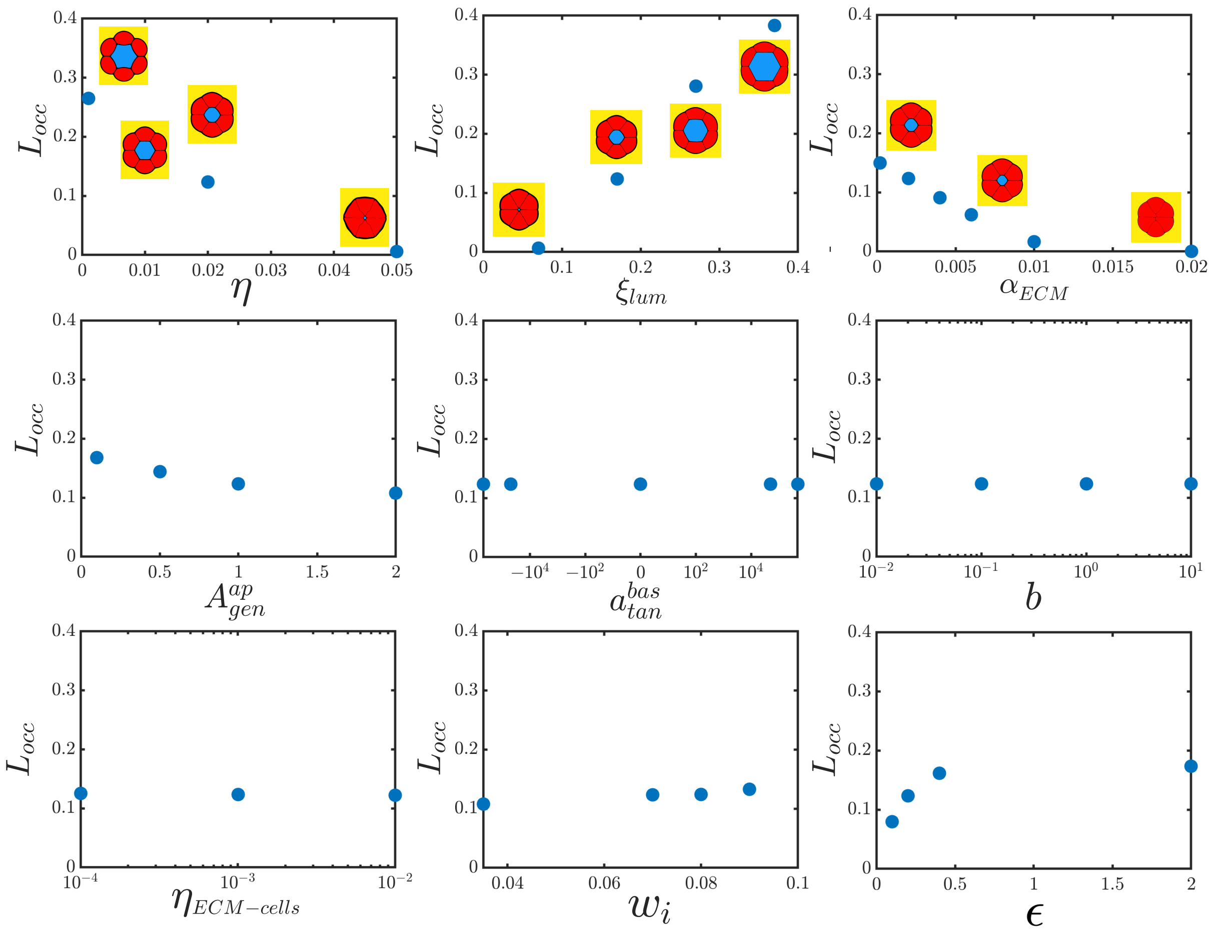

Supp. Fig. 7: ***Sensitivity analysis for lumen occupancy.*** *Sensitivity analysis. Lumen occupancy is measured as one parameter value is modified.*

We list in Supp. Table 2 the relevant parameters to our model. We identified a total of 41 parameters, accepting the redundancy, of which :

- 34 free parameters (in green cells of Supp. Table 2) and 31 relevant free parameters (because we can assume 3 units of length, time and force).
- 1 bounded parameters (blue cell of Supp. Table 2)
- 6 parameters estimated from experimental measurements (pink cells of Table 2)

The two rightmost columns of the Table 2 are related to sensitivity analysis of key relevant free parameters w.r.t two metrics for $\alpha = L_{\mathrm{ap}}/L_{\mathrm{bas}}$ and $L_{\mathrm{occ}}$. These values are measured by estimating the slope of plots of Supp. Fig. 6 and Supp. Fig. 7. This sensitivity analysis confirmed that varying cell-cell adhesion and lumen pressure was the most logical choice to find the steady-state of Fig. 3(e) where active gel parameters had been adjusted to experimental values.

| Parameter | Meaning | Num. value | Exp. value | Sensitivity | |
|---|---|---|---|---|---|
| | | | | $\alpha$ | $L_{occ}$ |
| $N\Delta x$ | Grid size | 12 | - | - | - |
| $\tau_\phi$, $\tau_\ell$, $\tau_c$ | Cell, lumen and ECM phase-field characteristic time | 1 | - | - | - |
| R | Cell radius | 1 | - | - | - |
| $V_\phi$ | Target cell surface | $\pi R^2$ | - | - | - |
| $\alpha_\phi$ | Cell phase-field growth rate (elastic spring constant) | 1 | - | - | - |
| $\beta_\phi$, $\beta_{\phi\ell}$, $\beta_{\phi c}$, $\beta_{c\ell}$ | Cell-cell, cell-lumen, cell-ECM, ECM-lumen surface exclusion | 1 | - | - | - |
| $\eta_\phi$ | Cell-cell adhesion | 0.02 $\in [1e^{-3}, 5e^{-2}]$ | - | -8.0 | -6.9 |
| $\eta_{\phi c}$ | Cell-ECM adhesion | 0.001 $\in [1e^{-4}, 1e^{-2}]$ | - | -1.2 | 0.3 |
| $\xi$ | Lumen pressure | 0.17 $\in [0.07, 0.37]$ | - | 1.8 | 1.3 |
| $\alpha_c$ | ECM elasticity | 0.002 $\in [2e^{-5}, 2e^{-2}]$ | - | -20 | -17 |
| $V_c$ | Target ECM surface | 144 | - | - | - |
| $\xi_c$ | ECM pressure | 0 | - | - | - |
| $A_{deg}^{apical/basal/lateral}$ | Degradation rate of the apical, basal, lateral active gel | Baseline: 100 Asymmetric case: Exp. determined | - | - | - |
| $\Gamma_{ap-bas}$, $\Gamma_{ap-lat}$ | Degradation rate ratios Apical/basal - Apical/lateral | $A_{deg}^{apical}/A_{deg}^{basal}$ $A_{deg}^{apical}/A_{deg}^{lateral}$ | 0.25 | - | - |
| $A_{gen}^{apical/basal/lateral}$ | Production rate of the apical, basal, lateral active gel | 1.0 $\in [0.1, 2.0]$ | - | $1.3e^{-2}$ | $-3.2e^{-2}$ |
| $\Lambda_{ap-bas}$, $\Lambda_{ap-lat}$ | Production rate ratios Apical/basal - Apical/lateral | $A_{gen}^{apical}/A_{gen}^{basal}$ $A_{gen}^{apical}/A_{gen}^{lateral}$ | 1.25 1.75 | - | - |
| $w = \sqrt{D}$ | Width of cortex | 0.07, $\in [0.035, 0.09]$ | - | $-7.4e^{-1}$ | $4.5e^{-1}$ |
| $\epsilon$ | Width of Dirac function | 0.2, $\in [0.1, 2.0]$ | - | $2.8e^{-2}$ | $4.9e^{-2}$ |
| $\gamma$ | Cortex - cytoplasm friction | 1 | - | - | - |
| $\eta$ | | 1 | - | - | - |
| $a_\perp^{apical/basal/lateral}$ $a_\parallel^{apical/basal/lateral}$ | Orthogonal/tangential active stress apical, basal, lateral | $\in [-5e^5, 5e^5]$ | - | 0.0 | 0.0 |
| $\Theta_\parallel^{ap-bas}$, $\Theta_\parallel^{ap-lat}$ | Tangential active stress ratios Apical/basal - Apical/lateral | $a_\parallel^{apical}/a_\parallel^{basal}$ $a_\parallel^{apical}/a_\parallel^{lateral}$ | 187.5 | - | - |
| b | Active gel saturation constant | 1, $\in [0.01, 10]$ | - | 0.0 | 0.0 |
| $f_{coup}$ | Coupling phase-field/active gel | 1 | - | - | - |

Supp. Table 2: ***Parameter table with numerical and experimental values as well as sensitivity analysis.*** *The sensitivity parameters are defined by the slope measured in Supp. Fig. 6 and Supp. Fig. 7 for the two key readouts $\alpha$ and $L_{occ}$. Green cells represent free parameters, blue cells bounded parameters and pink cells experimentally determined parameters.*

#### 4.5 Comparison between numerical simulation parameters and physical units

To compare experimental measurements with numerical simulations, we estimated numerical parameters with physical units. It is not straightforward to estimate cell cortical tension from our model parameters. To do so, we use equation (18) to derive that $A_{\text{gen}} \propto 1/T$, where $T$ is a characteristic timescale, $A_{\text{gen}} \propto \rho L/T$, where $\rho$ is a characteristic gel density, $L$ is a characteristic length scale (typically cell radius). Then, we use equations (21) and (23) to show that the cortical cell tension $\gamma_{\text{cell}}$ can be estimated by:

$$\gamma_{\text{cell}} \propto w_i \Pi \propto w_i a \rho^3 \propto w_i a \left( \frac{A_{\text{gen}}}{L A_{\text{deg}}} \right)^3$$

We evaluate $\gamma_{\text{cell}} \simeq 3.5 \cdot 10^{-2}$ with typical values used in simulations: $w_i = 0.07$, $a = 500000$, $A_{\text{gen}} = 1.0$, $A_{\text{deg}} = 100.0$ and $L = 1.0$.

Together with estimation of physical units following our previous work [41] where typical pressures are of the order of 1kPa, typical length are of the order of 10$\mu$m, $\gamma_{\text{cell}} \simeq 350$Pa.$\mu$m. Of note, the parameters used in our study compare to the parameters used in Ref. [41]. The main contribution to $\gamma_{\text{cell}}$ arises from the contractile and extensile stresses within the actomyosin cortex. Typical values of such stresses are of the order of 1000 Pa [73] with a typical cortical thickness of $\sim$ 200 nm [74]. It is important to state that even though simulation frameworks are slightly different between this present study and Ref. [41] because we implemented a cell cortical tension as well as ECM elasticity that were encoded differently or not at all present in their study, we both implemented competing forces of comparable strength. For instance, the order of magnitude of the ECM Young Modulus scales with the difference in lumen hydrostatic pressure ($\sim$ 100 Pa [75]) and cell-cell adhesion and cell surface tension parameters are comparable with each other [41].
